# A giant virus genome is densely packaged by stable nucleosomes within virions

**DOI:** 10.1101/2022.01.15.476465

**Authors:** Terri D. Bryson, Pablo De Ioannes, Marco Igor Valencia-Sánchez, Jorja G. Henikoff, Paul B. Talbert, Bernard La Scola, Karim-Jean Armache, Steven Henikoff

## Abstract

The two doublet histones of Marseillevirus are distantly related to the four eukaryotic core histones and wrap 121 basepairs of DNA to form remarkably similar nucleosomes. By permeabilizing Marseillevirus virions and performing genome-wide nuclease digestion, chemical cleavage and mass spectrometry assays, we find that the higher-order organization of Marseillevirus chromatin fundamentally differs from that of eukaryotes. Marseillevirus nucleosomes fully protect DNA within virions as closely abutted 121-bp DNA wrapped cores without linker DNA or phasing along genes. Likewise, we observed that a large fraction of the nucleosomes reconstituted onto multi-copy tandem repeats of a nucleosome positioning sequence are tightly packed. Dense promiscuous packing of fully wrapped nucleosomes rather than “beads-on-a-string” with genic punctuation represents a new mode of DNA packaging by histones. We suggest that doublet histones have evolved for viral genome protection and may resemble an early stage of histone differentiation leading to the eukaryotic octameric nucleosome.

## Introduction

The association of histones with DNA in the eukaryotic nucleus was known by the late 19^th^ century (Luck, 1965), but it was the revolutionary discovery of nucleosomes in the early 1970s (Hewish and Burgoyne, 1973; Kornberg and Thomas, 1974; Noll, 1974) that established the fundamental subunit structure of eukaryotic chromatin. Other nucleosome configurations were described for homotetrameric archaeal histones, which were found to wrap ∼60-bp of DNA (Pereira et al., 1997) and form higher-order “slinkies” (Mattiroli et al., 2017). Some archaeal histones are doublets with two histone fold domains, a differentiated form hypothesized to predate the evolution of eukaryotic nucleosomes (Malik and Henikoff, 2003). Although the DNA wrap of archaeal nucleosomes resembles that of eukaryotic nucleosomes, the histones are too dissimilar in sequence from the four core eukaryotic histones to identify correspondences with specific domains of histone doublets. However, sequencing of the giant Marseillevirus discovered in 2007 led to the realization that its genome encodes two histone doublets that are paired homologs of the four eukaryote core histones (Boyer et al., 2009). In Marseillevirus, Hβ-Hα is homologous to eukaryotic H2B and H2A, and Hδ-Hγ is homologous to H4 and H3. Related viruses of the family *Marseilleviridae* have since been discovered infecting *Acanthamoeba* species worldwide (Sahmi-Bounsiar et al., 2021). Phylogenetic analysis places Hα, Hβ, Hγ, Hδ as sister groups respectively to their H2A, H2B, H3 and H4 eukaryotic counterparts, consistent with divergence from the last eukaryotic common ancestor for all four histone folds (Erives, 2017).

We and others have previously used biochemical reconstitution and cryoEM imaging to solve the high-resolution structure of Marseillevirus nucleosomes, which show a striking resemblance to their eukaryotic counterparts (Liu et al., 2021; Valencia-Sanchez et al., 2021). Unlike octameric eukaryotic nucleosomes, which wrap 147 bp of DNA, tetrameric Marseillevirus nucleosomes wrap only 121 bp of DNA, despite being reconstituted on the Widom 601 artificial positioning sequence, selected for 147-bp wrapping of nucleosome cores. Because Marseillevirus doublets assemble into nucleosomes that are not fully wrapped by DNA, it was proposed that they are inherently unstable, perhaps to facilitate expression during early stages of infection or for gene regulation (Liu et al., 2021). However, without isolation of viral chromatin in its native form, the extent to which reconstituted Marseillevirus nucleosomes are representative of their conformation in virions is debatable (Vannini and Marazzi, 2021). By mapping the chromatin landscape of the viral genome we aimed to understand the functional and evolutionary basis for viral nucleosomal packaging.

Here we show that chromatin from Marseillevirus can be released from viral particles by breaching the capsid and permeabilizing the lipid membrane followed by either enzymatic or chemical cleavage. Using mass spectrometry, MNase-seq and MPE-seq, we confirmed that the 121-bp wrap observed by cryoEM for reconstituted nucleosomes is also observed for endogenous histone doublets in the virion. However, in contrast to eukaryotic nucleosomes, which are 147-bp particles separated by ∼50-bp linkers, 121-bp Marseillevirus nucleosomes are tightly packed within the virion. Unlike eukaryotic nucleosomes, which are depleted from transcription start sites and phased downstream, we observed no depletion or phasing at genes. To determine whether tight packaging seen in the virion is inherent to Marseillevirus histone doublets, we reconstituted histones on multi-copy arrays with a strong 147-bp positioning sequence. We found that most of the arrays were fully occupied by histone doublets and were resistant to chemical cleavage, although a fraction of the nucleosomes were wrapped to form 147-bp particles, suggesting that neighboring nucleosomes stabilize wrapping. Our findings indicate that Marseillevirus histone doublets have evolved for tight packing of 121-bp particles without sequence or genic preference, consistent with a viral packaging function, thus representing a previously unknown mode of genome packaging by histones.

## Results

### Release of Marseillevirus chromatin from viral particles by nuclease digestion

The capsids of giant viruses that infect amoeba are resistant to treatments that disrupt other viruses. Previous attempts to release chromatin intact from *Marseilleviridae* capsids were reported to have been unsuccessful (Liu et al., 2021), however by dialyzing in low pH conditions Schrad and co-workers (Schrad et al., 2020) had shown that giant Mimivirus capsids can be breached, although without releasing their contents. We followed their protocol to open the Marseillevirus capsid, then dialyzed into a neutral buffer for Micrococcal Nuclease (MNase) digestion. Capillary gel electrophoresis revealed a striking ladder of protected particles with mono- and di-nucleosomes dominating and discernable tri- and tetra-nucleosome peaks (**Figure 1a**). Under the same permeabilization and digestion conditions, Drosophila mono-nucleosomes dominate, with only a minor di-nucleosomal peak (**Figure 1b**).

**Figure 1:**
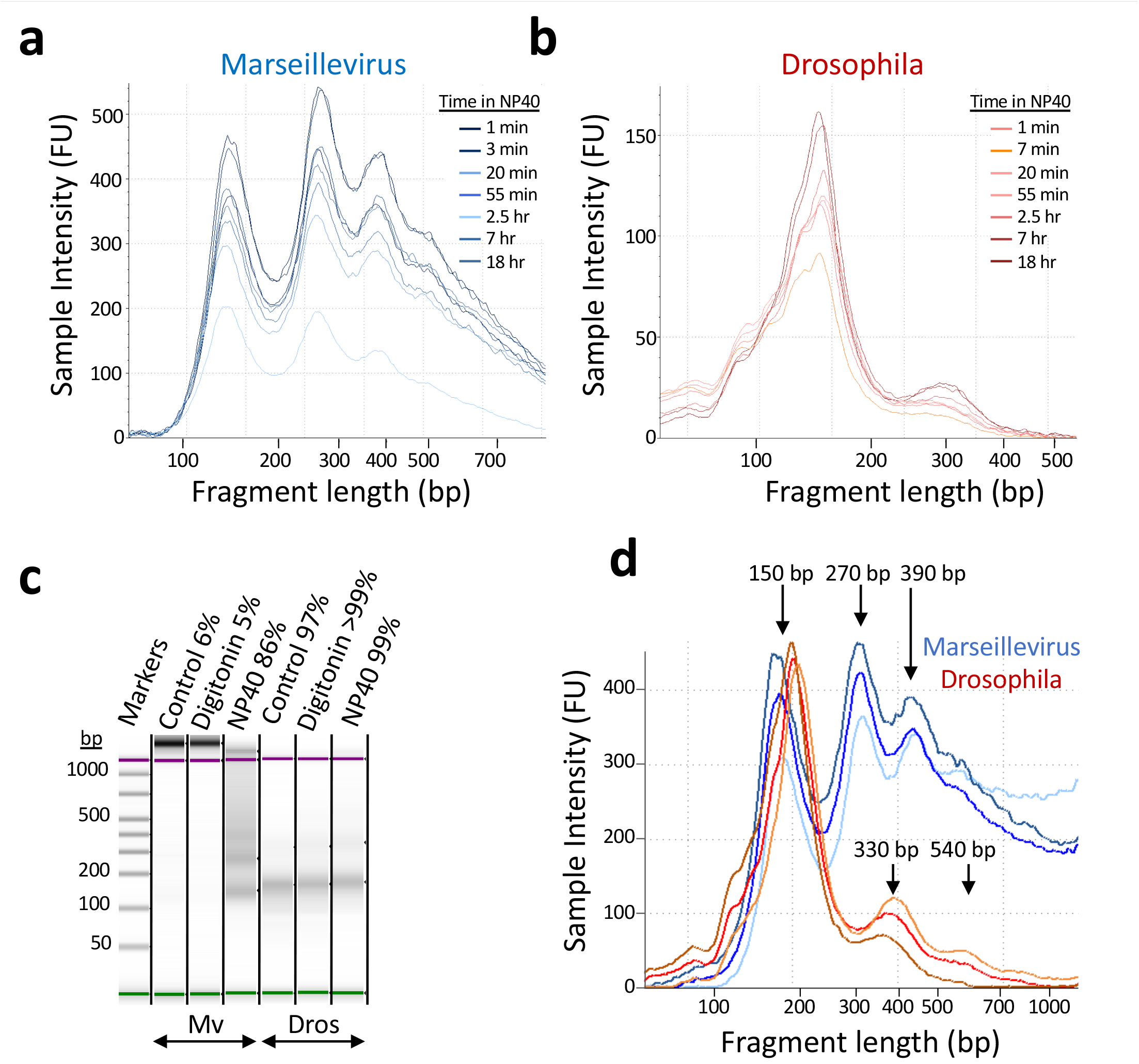
Permeabilization with NP-40 after capsid opening renders Marseillevirus chromatin immediately accessible to MNase digestion. (**a**) Electrophoretic mobility of Marseillevirus fragment lengths. NP40 completely permeabilizes pH2-treated and neutralized Marseillevirus particles within 1 minute to allow MNase digestion (10 U/million cells for 5 min at 37°C) with no significant changes after several hours. (**b**) Same as (a) for Drosophila chromatin after NP40 and MNase treatments under the same conditions. (**c**) Gel image of Marseillevirus (Mv) and Drosophila (Dros) chromatin extracted with no detergent (Control) or either Digitonin or NP-40. (**d**) Fragment length profiles of representative Marseillevirus and Drosophila NP40-permeabilized MNase digests. Estimated fragment sizes near ladder peaks are shown.

High yields of digested chromatin were obtained from samples treated with 0.5% NP-40 over a time-course ranging from 1 minute to 18 hours (**Figure 1a, S1a**). In contrast, 0.1% digitonin had no effect (**Figure 1c, S1b-c**). As NP-40 permeabilizes membranes by sequestering lipids, whereas digitonin permeabilizes cell membranes by displacing membrane sterols, our results suggest that the chromatin of *Marseilleviridae*, like that of giant *Mimiviridae* (Kuznetsov et al., 2010; Suzan-Monti et al., 2007; Xiao et al., 2009; Zauberman et al., 2008), is enclosed within an impermeable lipid membrane that would have protected it from low pH conditions, and subsequent permeabilization allowed for MNase to gain access to chromatin. Under our permeabilization and digestion conditions, 85-90% of the DNA in virions was recovered as intact MNase-protected particles (**Figure S1**).

To ascertain the protein composition of Marseillevirus chromatin, we centrifuged viral suspensions following NP40 permeabilization and MNase treatment, then extracted total protein from the pellet, supernatant and washes. We performed SDS-PAGE and excised bands from silver-stained gels as indicated (**Figure 2a**), including the ∼25 kDal bands corresponding to the predicted MW of both Hβ-Hα (25,949) and Hδ-Hγ (25,218) and neighboring bands. We also excised bands from total protein extracted from reconstituted chromatin using histones produced in *Escherichia coli* and from a blank gel lane. Protein samples were digested with trypsin and subjected to mass spectrometry followed by a comparison of peptide MWs to those predicted for trypsinized Marseillesvirus ORFs and likely contaminants. Only Hβ-Hα and Hδ-Hγ peptides were found at consistently high levels in both viral chromatin and reconstituted chromatin samples (**Figure 2b** and **Supplemental Item 1**). A distantly related variant of Hβ-Hα, Hζ-Hε (MW = 19,033), was detected in the chromatin fraction at a much lower level in the expected size range, consistent with previous mass spectrometry of viral extracts (Boyer et al., 2009). Taken together with the high DNA yield obtained from NP40-permeabilized and MNase-digested chromatin, our MS data confirm that the Hβ-Hα and Hδ-Hγ composition of viral chromatin matches that of the chromatin used for cryoEM (Valencia-Sanchez et al., 2021).

**Figure 2:**
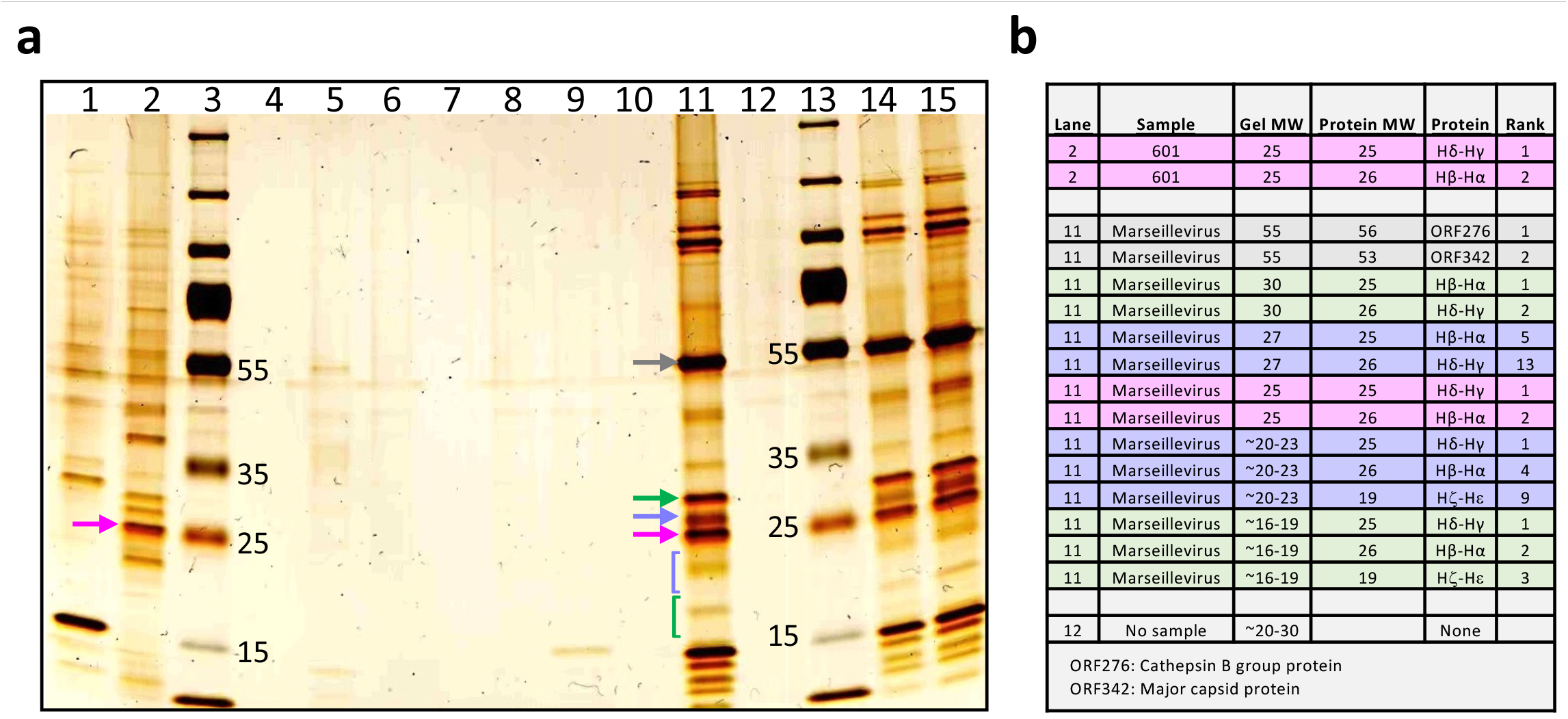
Proteomic analysis of viral protein extracts. (**a**) SDS-PAGE. Lanes are 1) MNase working stock; 2) Purified Marseillevirus histones reconstituted with 3x Widom 601 arrays after MNase digestion; 3) Marker proteins; 4) Blank; 5) Supernatant 1 (post-NP40 incubation, pre-MNase digestion); 6) Blank; 7) Wash (single wash in TM2); 8) No sample; 9) Supernatant 2 (post-MNase digestion); 10) Blank; 11) Pellet (post-MNase digestion); 12) No sample; 13) Marker proteins; 14) Mv-pH2 (crude lysate of unopened virus); 15) Virus suspension pH2 (crude lysate of breached virus). Gel slices (arrows and brackets) were excised and processed for MS. Doublet histones are ∼25 kDa (magenta arrows), capsid protein is ∼55 kDa (grey arrow) and the variant Hζ-Hε is ∼19 kDa. (**b**) MS summary. Predicted Marseillevirus proteins were rank-ordered by Sum PEP score (Supplementary Item 1) and only predicted proteins in the expected size range for the gel slice and predicted doublet histones were counted. Gel MW is estimated from the markers and Protein MW is predicted from the GenBank annotation (NC_013756.1, 457 open reading frames). Background colors correspond in order to the arrows and brackets in (a).

### Global analysis of the Marseillevirus genomic landscape

We performed paired-end DNA sequencing of Marseillevirus fragments and used Bowtie2 to align them to the Genbank T19 (later named *Marseillevirus marseillevirus*) genome assembly. This mapping revealed over-representation of fragments relative to the rest of the genome in two sharply defined regions (**Figure 2a**). A ∼20-kb region centered over Position 27,000 is ∼3-fold over-represented, and a ∼2-kb region centered over Position 318,400 is ∼10-fold over-represented similarly in all ten MNase datasets regardless of digestion level. To determine whether these striking over-representation features were present in the initial sample from 2007 used to assemble the original T19 map, we mapped the primary T19 reads from the original fastq files, and observed that over-representation at Position 27,000 was already conspicuously present in this sample (**Figure 3a**, left, top track).

**Figure 3:**
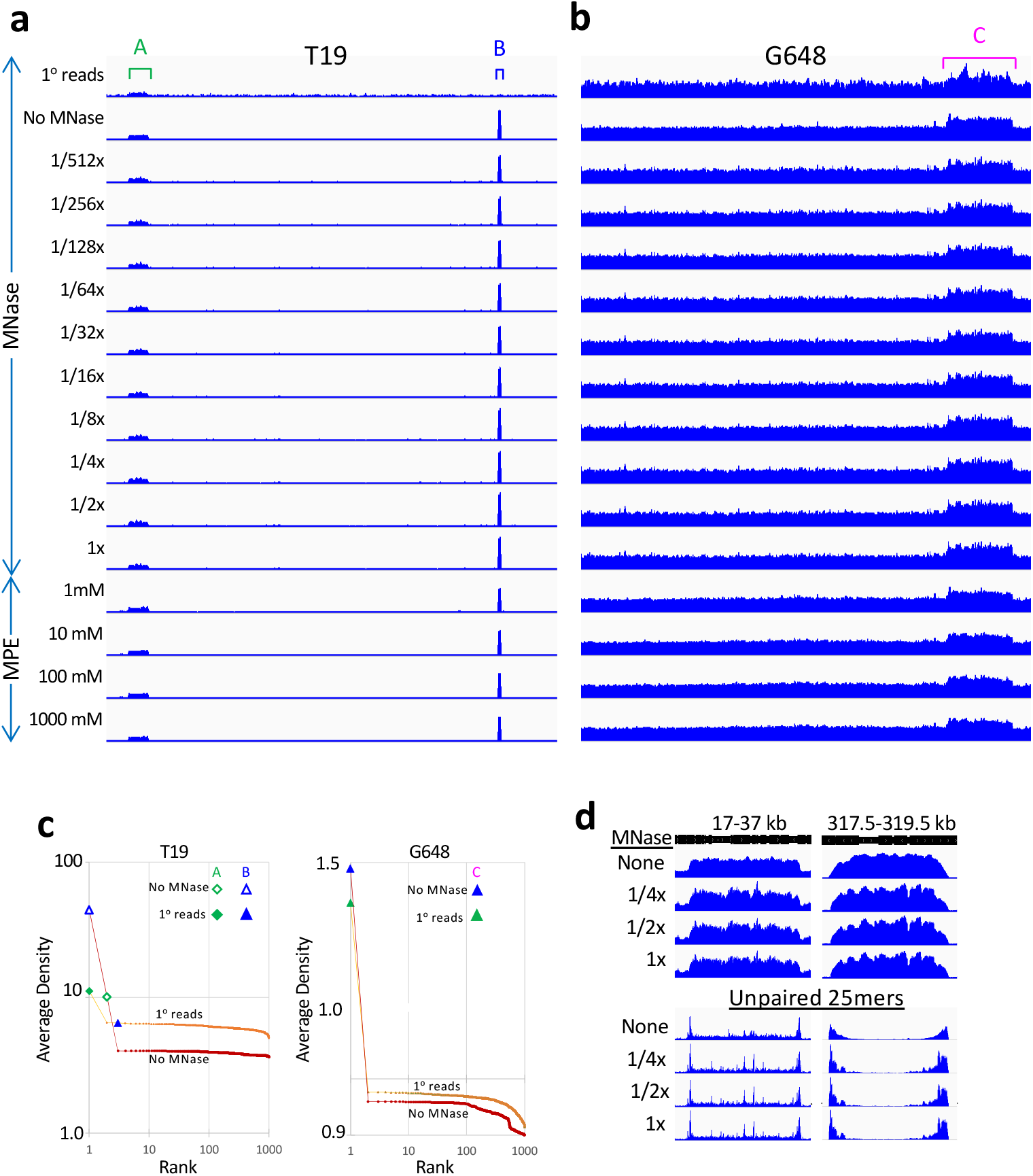
Regional over-representation of DNA during culturing of Marseillevirus. (**a**) IGV screenshots reveal over-representation of specific regions detected using raw reads from the primary *Marseillevirus marseillevirus* (T19) assembly (8), sequencing without MNase treatment (No MNase), an MNase-seq digestion series (where 1x = 10 U/million cells, 1/2x = 5 U/million cells, and so on) and an MPE-seq digestion series (1 μM, 10 μM, 100 μM and 1000 μM H_2_O_2_). (**b**) Same as (a) for G648 using the published assembly (8). (**c**) Cumulative plot showing the rank of each over-represented region among random samples of the span from regions of the rest of the genome, where Rank 1 is the most over-represented sample. We excluded 1 kb on either side of the regions and at either end of the linearized genome assembly of the circular genome. (**d**) To identify novel junctions at the borders of over-represented regions we compared maps of total fragment occupancy to maps of 25-mers, where only a single 25-mer fragment end was mapped (Unpaired). The Y-axis is autoscaled within groups. A novel junction will be recognized because any fragment spanning the junction without overlap will not be mapped, as the 25-mer ends will be oriented 3’-to-5’ relative to one another, rather than 5’-to-3’.

To confirm the validity of the Genbank assembly of the T19 circular genome, we performed *de novo* genome assembly for a representative T19 sample. We ran the SPAdes program on the 50-bp end reads, which provided >2000x genomic coverage per sample. We obtained >99.9% coverage by the 8 largest contigs, where regions of over-representation are spanned in our assembly (**Table S1**). This demonstrates that the over-represented regions seen in T19 and G648 are inherent to the genome used for profiling chromatin, and do not reflect a mis-assembly artifact.

To quantify over-representation, we randomly sampled fragments of similar lengths to each over-represented feature from the rest of the genome 1000 times and plotted the abundance of each sample on a log-log cumulative plot for the raw reads from the original 2007 virus culture and the No-MNase paired-end reads from the 2020 virus culture used in this study (**Figure 3c**, left panel). This revealed that the 20-kb region over Position 27,000 was ∼2-fold over-represented in the original 2007 viral culture and ∼2.7-fold over-represented in the 2020 culture relative to their respective genome-wide median abundances. Likewise, the 2-kb region over Position 318,400 was ∼1.1-fold over-represented in the original 2007 viral culture and ∼10-fold over-represented in the 2020 culture. Although the magnitude of over-representation of the 2-kb region in the original 2007 viral culture is small, it is ranked between #1 and #2 of the 1000 randomly sampled regions, and so likely represents incipient over-representation of some virions in that culture. It is evident that over-representation of the 20-kb region continued to be maintained at 2-3-fold over-abundance during successive passages in *Acanthamoeba polyphaga* culture at similar levels within the cultured Marseillevirus populations, while the 2-kb region expanded ∼12-fold during successive passages in culture.

We next asked if regional over-representation and expansion during culturing is a general feature of the Marseilleviridae (Sahmi-Bounsiar et al., 2021). G648 is a much more recent isolate and therefore has been maintained for a shorter time in culture than T19. For G648 we observed a different region of over-representation, in which a ∼50 kb segment approximately centered over Position 319,000 is present in all ten samples in the MNase concentration series (**Figure 3b**). Although the primary reads from the fastq files used for the original assembly of G648 were relatively sparse, quantitative sampling analysis showed that this ∼50-kb region is also strongly over-represented, with a 1.9-fold excess, compared to a 2.6-fold excess for the No MNase sample (**Figure 3c**, right panel). As for T19, *de novo* assembly yielded >99.9% coverage by the 8 largest contigs, confirming the validity of the original assembly (**Table S1**).

Mapping of unpaired fragments from paired-end sequencing libraries shows them to be in large excess immediately outside of the over-represented regions as expected for the presence of novel tandem repeat junctions (**Figure 3d**). Our observations of different over-represented regions in two Marseillevirus isolates and increases in the abundance of over-represented regions in T19 during passage indicate that regional over-representation occurs during culturing of Marseillevirus in *A. polyphaga*. Analysis of long repeat-spanning fragments indicated that regional over-representation is accounted for by intrachromosomal tandem repeats rather than extrachromosomal circles (**Figure S2**).

### The Marseillevirus genome is densely packaged by 121-bp nucleosomes without linkers

Marseillevirus mono-nucleosomal DNA fragments released by MNase are smaller than those of Drosophila, and show much less spacing between nucleosomes (**Figure 1d**). Comparison of a Marseillevirus ladder to a lightly digested Drosophila ladder revealed an average difference of ∼55 bp for di-nucleosomes (two nucleosomes separated by one linker) and ∼120 bp for tri-nucleosomes (three nucleosomes separated by two linkers). Given that the average eukaryotic nucleosome repeat length is ∼200 bp and a nucleosome wraps 147 bp, our MNase digestion results suggest that there are no linkers separating Marseillevirus nucleosomes, but rather that they are abutted against one another (**Figure S3a**). Such tight packing of Marseillevirus nucleosomes might explain why they remain insoluble even after MNase digestion, in contrast to Drosophila nucleosomes, which are quantitatively released by MNase into solution (**Figure 1c, Figure S1**), as if tight packaging within the permeabilized virion prevents release of mono-nucleosomes.

At the highest MNase digestion levels, control Drosophila nucleosomes showed a dominant 147-bp peak, but also a smear of sub-nucleosome-sized digestion products (**Figure 4a**). Using the highest digestion level for Marseillevirus we observed a dominant 121-bp MNase-protected peak, a small ∼70-bp peak, but no smear, confirming cryoEM observations of 121-bp wrapping of reconstituted nucleosome cores (Liu et al., 2021; Valencia-Sanchez et al., 2021). These observations indicate that Marseillevirus nucleosomes are less sensitive to intranucleosomal cleavages than are eukaryotic nucleosomes.

**Figure 4:**
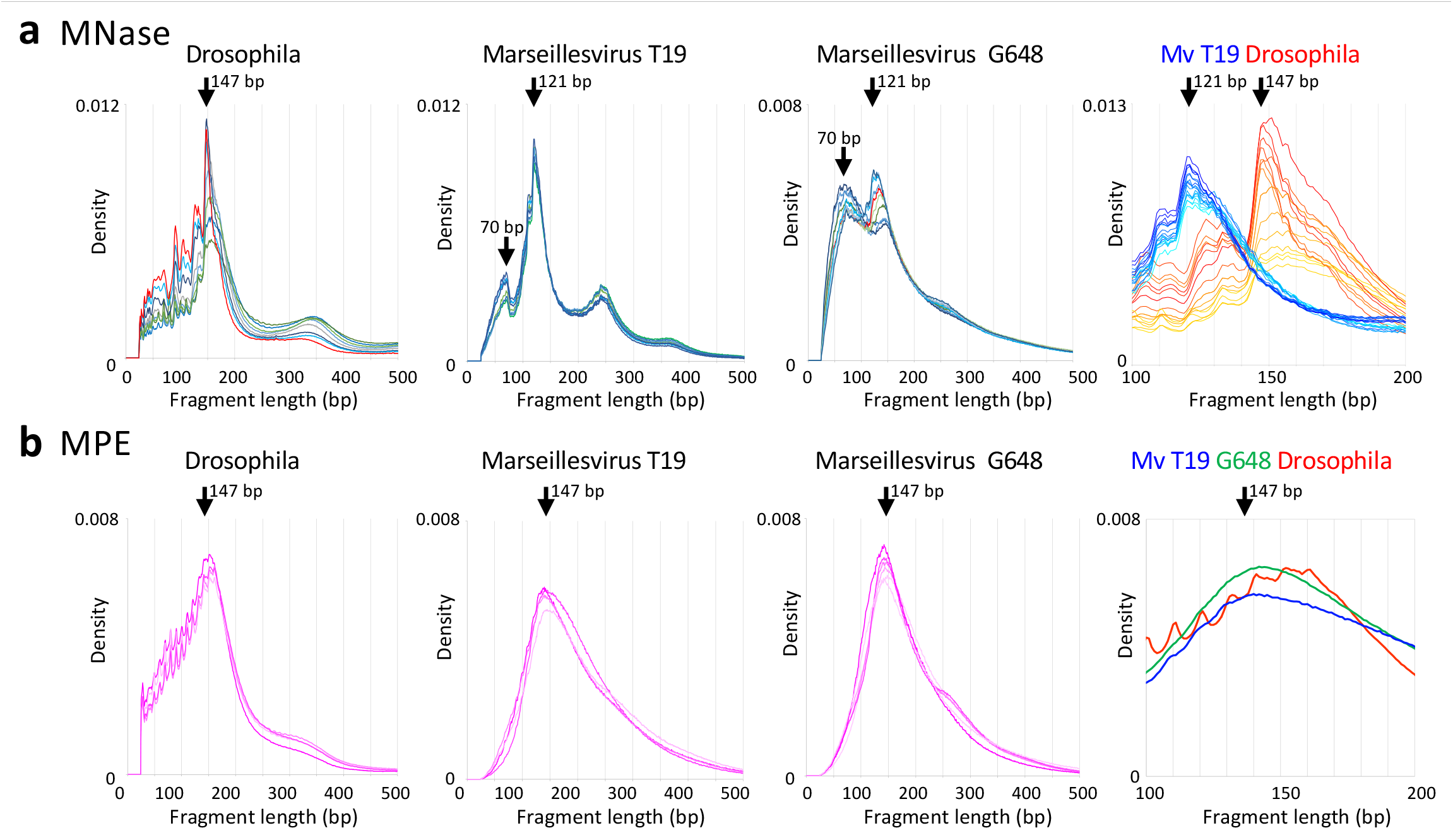
MNase- and MPE-digested Marseillevirus chromatin yields tightly packed nucleosome-sized DNA fragments. MNase-seq and MPE-seq were performed on Drosophila melanogaster S2 cells and on viral particles after pH 2 treatment and neutralization, followed by Illumina DNA library preparation and PE25 paired-end sequencing. (**a**) Drosophila (9 samples), Marseillevirus T19 extracted with NP40 (8 samples) and Marseillevirus G648 (10 samples) extracted without NP40. For MNase-seq, Drosophila and Marseillevirus G648 samples were digested for 5 min in a concentration series starting at 1x (red curve) and decreasing to 1/512x. Marseillevirus T19 samples were extracted for different times in NP40 and digested at the highest (1x) level used for Drosophila and Marseillevirus G648. Graphs on right show a comparison between Drosophila (14 samples) and Marseillevirus T19 (17 samples) on an expanded scale. (**b**) Same as (a) for MPE-cleaved samples, Drosophila (4 samples), Marseillevirus T19 (6 samples) and Marseillevirus G648 (4 samples). Curves for individual samples processed in parallel are superimposed to illustrate the degree of variation between replicate samples digested under different conditions used in this study.

### MPE-seq confirms the dense packing of Marseillevirus nucleosomes

MNase is an endo/exo-nuclease that is known to preferentially digest AT-rich regions (Chung et al., 2010; McGhee and Felsenfeld, 1983). Consistent with these observations, we found that ∼90% of the cleavage sites occur between A/T base pairs (**Figure S4**) and long AT-rich regions are preferentially digested (**Figure S5**). Therefore, we wondered whether the unexpected ∼70 bp peak (**Figure 4a**) resulted from aggressive MNase digestion of AT-rich regions within nucleosomes. To eliminate this potential artifact, we used a small-molecule DNA cleavage reagent, methidiumpropyl-EDTA-Fe(II) (MPE) which hydrolyzes DNA phosphodiester bonds, where H_2_O_2_ provides reactive oxygen for MPE-Fe(II)-catalyzed DNA cleavage (Cartwright et al., 1983). MPE-seq is performed similarly to MNase-seq, but without exonuclease activity and without sequence bias (Ishii et al., 2015) (**Figure S5**). When we treated Drosophila nuclei and permeabilized Marseillevirus particles with MPE-Fe(II), we observed mostly mononucleosome-sized particles for Drosophila, with a ∼150-bp peak and periodic 10-bp internal cleavages (**Figure 4b**, leftmost panel). MPE-seq of Marseillevirus chromatin from both T19 and G648 also revealed an ∼150-bp peak with close concordance between samples, and no 70-bp peak. Whereas MNase digestion resulted in a 26-bp fragment size difference between Marseillevirus and Drosophila nucleosomes (**Figure 4a** right panel), MPE cleavage resulted in fragments of nearly the same size (**Figure 4b**, right panel). These consistent discrepancies between MNase- and MPE-generated fragments indicate that the aggressive endo/exonucleolytic activity of MNase and its preference for AT-rich DNA is responsible for the ∼70-bp fragment peaks in T19 and G648. Although the genomes of both Marseillevirus isolates have the same ∼44% GC-content as Drosophila, they have characteristic A/T homopolymeric runs in promoter regions (Oliveira et al., 2017) that might contribute to artifactual internal MNase cleavages seen for Marseillevirus nucleosomes (**Figure S6**). The lack of periodic 10-bp internal cleavages with MPE digestion demonstrates that Marseillevirus chromatin is highly refractory to intranucleosomal cleavages (**Figure 4b**, rightmost panel, **Figure S3a**).

To examine the genome-wide organization of Marseillevirus nucleosomes, we separated fragments by size in 100-bp intervals and displayed a representative 10-kb region of the T19 and G648 genomes for MNase- and MPE-generated replicate samples. Typical Drosophila chromatin profiles showed patches of positioned nucleosomes for 101-200 bp fragments, but little if any consistent positioning between replicates for subnucleosomal 1-100 bp or supranucleosomal 201-300 bp fragments for both MNase-seq (**Figure 5a**) and MPE-seq (**Figure 5b**). Although we observed patches of positioning for Marseillevirus T19 and G648, this was seen for all three size classes, but only for MNase-seq, with little if any distinctness in the chromatin landscape for MPE-seq. We also examined these representative 10-kb regions for evidence of nucleosome phasing by performing autocorrelation analysis (**Figure 5c**). This revealed consistent periodicities for both the MNase-seq and MPE-seq Drosophila data, most prominently for the 101-200 subset, as expected for nucleosome phasing. In contrast, the slight periodicities we observed for Marseillevirus T19 and G648 were mostly inconsistent between MNase-seq and MPE-seq. This lack of consistent phasing in Marseillevirus T19 and G648 nucleosomes suggests that they are densely packed without intervening linkers characteristic of eukaryotic nucleosomes.

**Figure 5:**
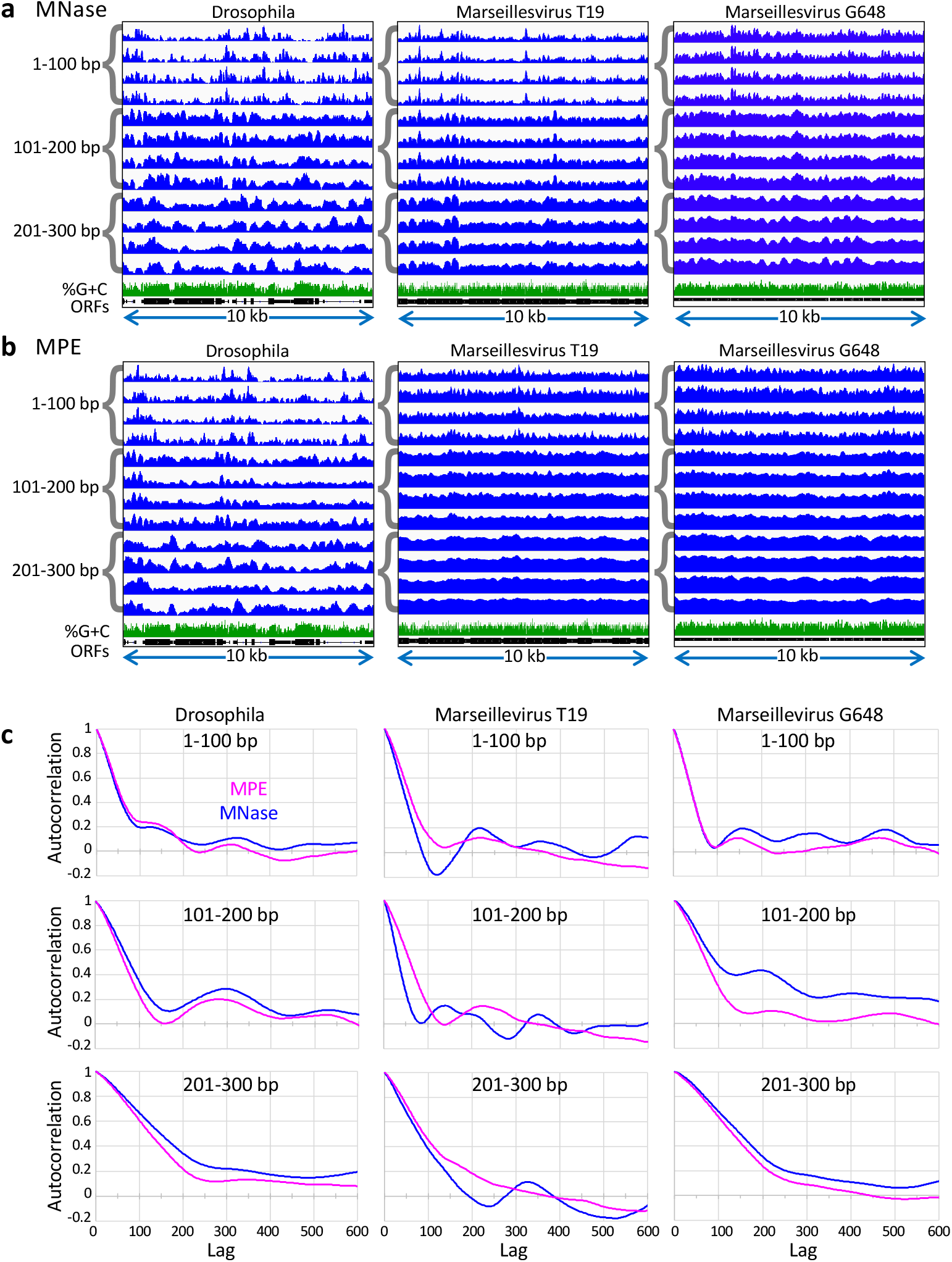
Marseillevirus chromatin is periodic without linker DNA. Drosophila and Marseillevirus T19 and G648 MPE-seq and MNase-seq library sequences aligned to their respective genomes, and four MNase samples (**a**: top to bottom: 6.25, 12.5, 25 and 50 mU/reaction) and four MPE samples (**b**: top to bottom: 1 μM, 10 μM, 100 μM and 1000 μM H_2_O_2_) were chosen for comparison. For Drosophila, the 10-kb region around a bidirectional housekeeping promoter pair (His2Av and ball, chr3R:26,862,501-26,872,500) was selected for display. For T19 an arbitrarily chosen representative 10-kb region was selected (53,001-63,000) and the region with the same coordinates in G648 (but not orthologous) was also selected. Fragment size classes represent subnucleosomes (1-100 bp), nucleosomes (101-200 bp) and mixtures of mono- and di-nucleosomes (201-300 bp). Nearly uniform occupancy is seen for MPE-generated Marseillevirus nucleosomes, whereas under the same conditions Drosophila nucleosomes display regions of conspicuous phasing characteristic of nucleosomes separated by linker regions. Tracks are group-autoscaled within sets of four. Percent G+C (green) and open reading frames (ORFs, black boxes) are plotted at 10-bp resolution for each 10-kb span. (**c**) Autocorrelation illustrates periodicities over representative regions. To sensitively detect nucleosome phasing, we plotted autocorrelations over the first 600 bp of each representative region shown above for the three fragment size classes as indicated using MNase-seq and MPE-seq data. Autocorrelations in 1-bp lag steps over a 5’-aligned 10-kb span are plotted for MNase-seq (blue) and MPE-seq (magenta).

### The Marseillevirus chromatin landscape lacks genic differentiation

We wondered whether the minor regularities seen in the MNase-digested Marseillevirus nucleosome landscapes and by autocorrelation analysis (**Figure S6**) correspond to genic regions, which are annotated as Open Reading Frames (ORFs) and are separated by very short spans that are usually AT-rich (Oliveira et al., 2017). To investigate this possibility, we aligned all 191 Open Reading Frames (ORFs) that are ≥600 bp at their 5’ ends and averaged each nucleotide position over gene bodies. For comparison we chose the first 191 ORFs ≥600 bp from the chronological list of Drosophila ORFs. Alignment of MNase-generated 101-200 bp fragments to the 191 Drosophila ORFs revealed the characteristic translational phasing pattern, with a prominent +1 nucleosomal peak just downstream of the ORF 5’ end and phased +2 and +3 nucleosomes with reduced occupancy for all digestion levels over a 128-fold range (**Figure 6a**). MPE-generated fragments showed the same 5’-aligned translational phasing patterns for 101-200 bp fragments, confirming that maps produced using MPE and MNase are approximately concordant for control eukaryotic nucleosomes. In contrast, MNase-generated Marseillevirus T19 and G648 101-200 bp fragments displayed little if any 5’ phasing when plotted on the same scale (**Figure 6a**), although when the scale was expanded a positioned +1 nucleosome was observed. However, no nucleosome positioning was seen for MPE-generated T19 and G648 5’-aligned ORFs (**Figure 6a, c**), indicating that the minor +1 nucleosome positioning that was seen with MNase is likely attributable to its aggressive endo/exonuclease activity (**Figure S3a**). The lack of 5’ phasing, a universal characteristic of eukaryotic genes, indicates that the packaged Marseillevirus genome is likely to be transcriptionally inactive upon release from the capsid during infection.

**Figure 6:**
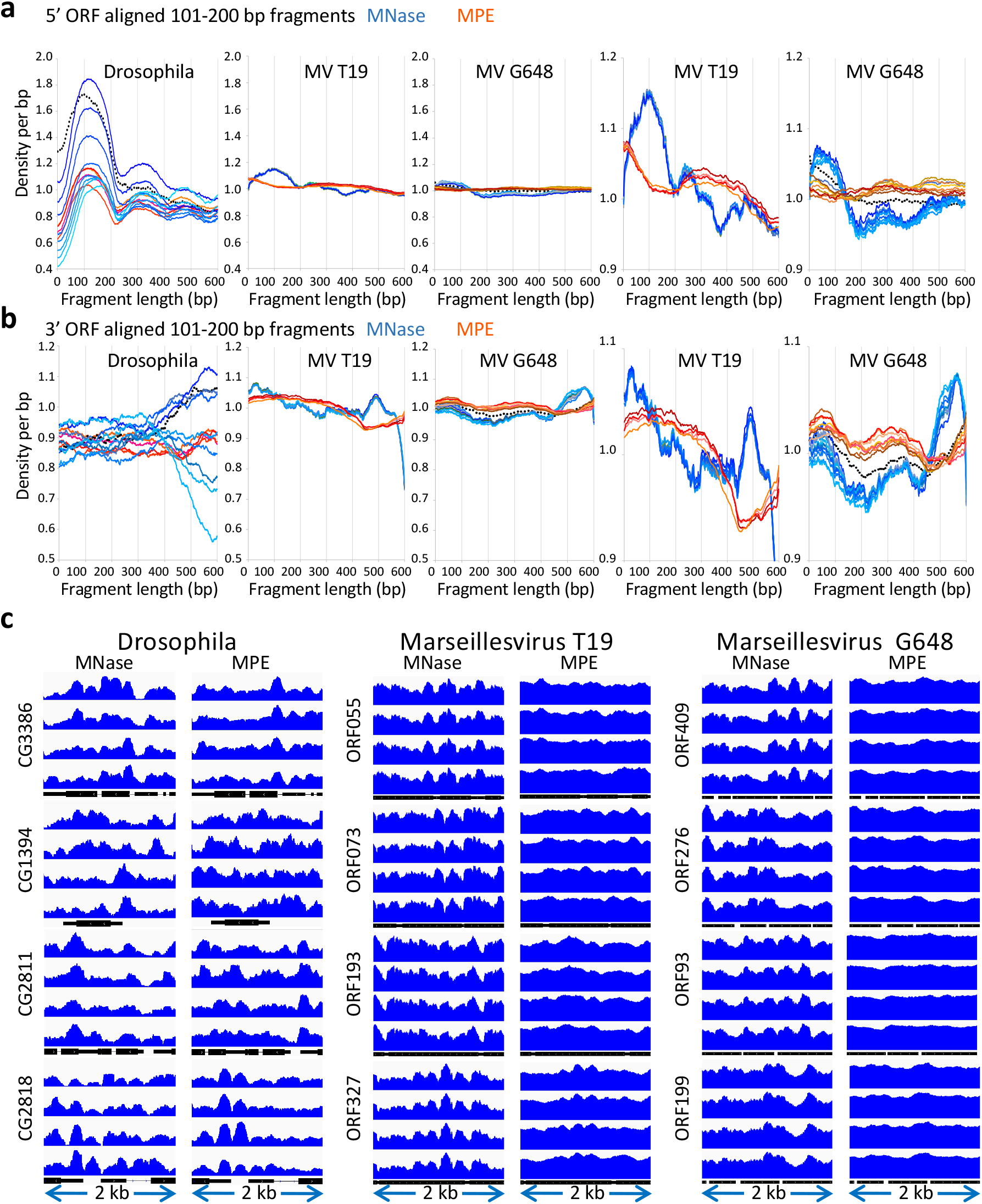
Phasing observed by MNase-seq for Marseillevirus is not observed by MPE-seq. All 191 Marseillevirus ORFs ≥600 bp were aligned (**a**) at their 5’ ends and (**b**) at the stop codon at their 3’ ends. Normalized counts are plotted at 1-bp resolution over the 600 bp span. Left to right: Drosophila S2 cells (8 MNase and 4 MPE samples), Marseillevirus T19 (8 MNase and 4 MPE samples) and G648 (10 MNase and 8 MPE samples) plotted on the same scale, and T19 and G648 plotted on a vertically expanded scale to display the differences between MPE and MNase digestions. Curves with bluish colors are from MNase-seq for different samples digested in parallel, and curves with reddish colors are for MPE-seq samples. Dotted line is for a no MNase control. (**c**) ORFs showing high variability based on autocorrelation (Figure S5) downstream of the 5’ end for the 101-200 kb size class were displayed within 2-kb regions for four technical replicates each. For MNase-seq, samples were digested in (top to bottom) 50, 25, 12.5, 6.25 mU MNase/reaction. For MPE-seq, samples were digested using (top to bottom) 1 μM, 10 μM, 100 μM and 1000 μM H_2_O_2_. In contrast to Drosophila nucleosomes, which show phasing of 101-200 bp fragments resulting from either MNase or MPE digestion, Marseillevirus nucleosomes are not detectably phased when assayed by MPE-seq, and MNase-seq shows phasing irrespective of fragment size, indicating internal cleavages (Figure S4). Tracks are group-autoscaled within sets of four.

Alignment of the 191 ORFs at the stop codon of their 3’ ends showed extreme sensitivity of Drosophila nucleosomes to MNase levels not seen for MPE-generated fragments (**Figure 6b**), which is consistent with partial unwrapping of AT-rich ORF 3’-end DNA from nucleosome cores and sensitivity to MNase exonucleolytic activity. A similar MNase sensitivity and MPE insensitivity was seen for T19 and G648 nucleosomes at the very 3’ ends of ORFs (**Figure 6b**), which is likewise attributable to AT-rich regions at Marseillevirus 3’ ends. However, unlike chromatin at the 3’ ends of Drosophila ORFs, both T19 and G648 chromatin displayed an average peak of MNase resistance just upstream of the 3’ end. As this peak was absent from MPE-digested average profiles, we attribute it to internal MNase cleavage of a subset of nucleosomes that are relatively excluded from neighboring AT-rich regions rather than to 3’ nucleosome phasing. The lack of any chromatin accessibility features punctuating genic regions implies that Marseillevirus nucleosomes have evolved exclusively for packaging within the virion.

### Dense packing of reconstituted Marseillevirus chromatin on Widom 601 arrays

We wondered whether previous reconstitutions of Marseillevirus nucleosomes on single 601 sequences (Liu et al., 2021; Valencia-Sanchez et al., 2021) failed to show fuller wrapping because of the lack of neighboring nucleosomes, which we find are closely abutted in virions. Accordingly, we performed MPE and MNase digestion on native and cross-linked reconstituted Marseillevirus chromatin that had been assembled at 60 μg/ml or 300 μg/ml concentrations onto a three-copy 601 array and onto a 12-copy 601 array. Following DNA extraction, we prepared sequencing libraries and performed paired-end sequencing, aligning fragments from native or cross-linked 3-copy and 12-copy chromatin digests to the 3-copy 601 array. For the 3-copy native chromatin, MPE-seq revealed similar chromatin landscapes for the 60 μg/ml and 300 μg/ml samples, with higher occupancy over the three 147-bp 601 positioning sequences than over the intervening 40-bp linkers, although not as high as for Xenopus control core histones assembled on the same 3-copy array (**Figure 7a**, left). The transitions between 601 and linker sequence were sharply defined, indicating precise positioning of 147-bp 601 particles over the 601 sequence. A much smaller fraction of 147-bp 601 particles was observed for MPE digestion of cross-linked 3-copy chromatin (**Figure 7a**, right).

**Figure 7:**
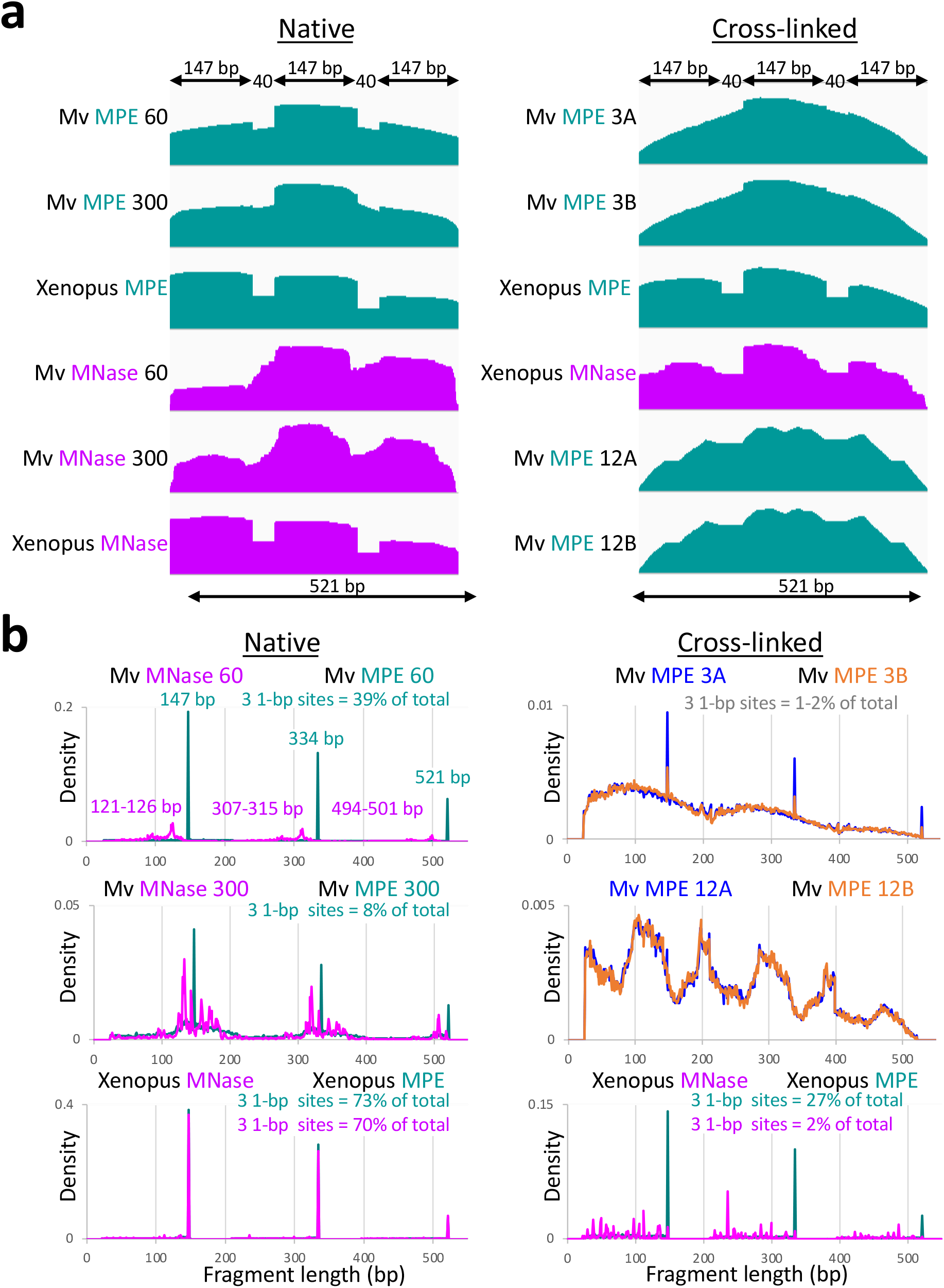
Assembly of Marseillevirus histones onto Widom 601 arrays produces 147-bp nucleosomes. (**a**) Three-copy (521 bp) or 12-copy (2084 bp) Widom 601 arrays and 60 μg/ml or 300 μg/ml of Marseillevirus histones and control Xenopus core histones were assembled into nucleosomes. Native (left) and cross-linked (right) assemblies were subjected to MPE-seq (cyan) and MNase-seq (magenta). The higher abundance of fragments towards the middle of each array is a consequence of mapping to a linear fragment, where fragments that span an end can only align with internal copies. MPE 12A and MPE12B were assembled onto the 12-copy array, cross-linked, digested with MPE and aligned to the 3-copy array. Profiles were group autoscaled. (**b**) Length distributions of fragments produced by MPE from the 3-copy array reconstituted samples show that up to 39% of the total occupancy is accounted for by precise cleavages on either or both ends of a 147-bp particle.

To quantify the relative abundance of cleavages precisely at 147-bp 601 particle ends, we plotted the length distributions for each 3-copy sample (**Figure 7b**). In each case, MPE digestions of 3-copy array chromatin resulted in 1-bp wide fragment length peaks at 147 bp, 334 bp and 521 bp. As the entire 3-copy array is 521 bp and 334 bp is exactly the size expected for a 601-linker-601 spanning fragment, it is evident that MPE endonucleolytically digests to completion without detectable encroachment into the nucleosome-wrapped particles. Nineteen percent of the nucleosomes in the 60 μg/ml sample were 147 bp and precisely phased over the 601 positioning sequence, and 147 bp, 334 bp and 521 bp fragment ends accounted for 39% of the total (**Figure S3b**). By comparison, control Xenopus nucleosomes assembled on the same 3-copy 601 array and subjected to MPE-seq yielded 38% 147-bp particles, and 147 bp, 334 bp and 521 bp fragment ends accounted for 73% of the total. This confirms that the inherent tendency of adjacent Marseillevirus nucleosomes to interact and partially overcome DNA sequence-directed positioning more effectively than that observed for eukaryotic nucleosomes. High histone occupancy over the linkers between 601 positioning sequences contrasts with the 121-bp wrap seen for Marseillevirus nucleosomes reconstituted onto single 601 sequences (Liu et al., 2021; Valencia-Sanchez et al., 2021) and suggests that closely abutted Marseillevirus nucleosomes stabilize one another and prevent unwrapping (**Figure S3a)**.

After cross-linking, only ∼1% of 3-copy Marseillevirus chromatin arrays corresponded to 147 bp particles, and 147 bp, 334 bp and 521 bp fragment ends accounted for only ∼2% of the total, superimposed over a broad distribution of fragment lengths (**Figure 7b** and **S3b**). For 12-copy cross-linked reconstituted arrays, MPE-seq resulted in the total absence of 1-bp peaks. By comparison, control Xenopus nucleosomes reconstituted on 3-copy arrays yielded 14% 147-bp particles, and 147 bp, 334 bp and 521 bp fragment ends accounted for 26% of the total (**Figure S3b**). This indicates that a larger fraction of the MPE cleavages are uniformly distributed on reconsistuted Marseillevirus chromatin than on reconstituted eukaryotic chromatin.

In contrast to the results with MPE, MNase digestion of the same reconstituted chromatin showed a distribution of fragment lengths ∼20-25 bp smaller than the three discrete fragment lengths produced by MPE digestion of Marseillevirus chromatin. Most notably, MNase digestion of native Marseillevirus chromatin produced fewer than 1% precisely positioned cleavages and resulted in a rough profile, whereas for Xenopus nucleosomes assembled on the same 3-copy Widom 601 array, 70% of the cleavages were precisely positioned and resulted in a clean sawtooth pattern (**Figure 7a** and **S3b**). Reduced nucleosome positioning on 601 arrays measured by both MPE-seq and MNase-seq distinguishes Marseillevirus doublet histones from eukaryotic core histones and recapitulates the situation *in virio*.

## Discussion

We have shown that Marseillevirus nucleosomes can be recovered intact from virions and used to elucidate nucleosome organizational features using mass spectrometry, MNase-seq and MPE-seq. These methods reveal particles that differ from eukaryotic nucleosomes in being refractory to internal cleavages and tightly packed into a landscape without linker DNA or phasing around genes. Taken together, our findings reveal a dense chromatin landscape that may have evolved to maximize protection of viral DNA for survival during infection in amoeba cytoplasm. This mode of chromatin organization differs drastically from that of eukaryotes, where nucleosomes not only protect DNA, but also have evolved for gene regulation by limiting access to regulatory elements (Kornberg and Lorch, 2020). Given the close structural superimposition of the Marseillevirus nucleosome with the eukaryotic nucleosome (Liu et al., 2021; Valencia-Sanchez et al., 2021), our finding that Marseillevirus chromatin lacks linkers or genic punctuation is especially remarkable. Marseillevirus chromatin is also unlike that of well-studied archaeal chromatin, in which 60-bp single-wrapped units are thought to form long “slinkies” that dynamically open and close (Bowerman et al., 2021; Mattiroli et al., 2017). Rather, tight packing without linkers implies an inherently stiff fiber. Therefore, Marseillevirus nucleosomes represent a previously unknown mode of chromatin packaging.

In addition to traditional MNase-seq, we applied MPE-seq, a chemical cleavage mapping method with high penetrability and without sequence bias (Ishii et al., 2015). Because MPE-seq lacks “nibbling” activity, it revealed fully wrapped 147-bp reconstituted particles on Widom 601 DNA. This artificial nucleosome positioning sequence was based on SELEX selection from chemically synthesized random DNA sequences using eukaryotic histone cores (Lowary and Widom, 1998). Therefore, our results using reconstituted 601 arrays imply that Marseillesvirus nucleosomes follow the same rules for nucleosome wrapping as eukaryotic nucleosomes, but doublet histones have evolved to wrap less DNA to facilitate dense nucleosome packing without intervening linkers. The fact that wrapping to the edges of the 147-bp Widom positioning sequence on 3-copy arrays accounts for only ∼40% of the total and just 2% when cross-linked implies that the large majority of MPE-generated cleavages result from dense nucleosome packing without sequence preference on arrays just as we observed in virions.

The lack of linkers or genic punctuation of chromatin in virions raises questions as to how the chromatin landscape becomes accessible following infection, when the virion is transformed into a cytoplasmic “viral factory” within *Acanthamoeba* cytoplasm (Liu et al., 2021; Suzan-Monti et al., 2007). In addition to the abundant doublet Hβ-Hα and Hδ-Hγ histones that Marseillevirus encodes on divergent transcription units, a separately encoded histone doublet variant, Hζ-Hε, is distantly related to Hβ-Hα and is present at very low levels in the chromatin fraction of the virion (**Figure 2b**). It is possible that Hζ-Hε acts as a replacement variant, analogous to the eukaryotic histone variant H2A.Z, which replaces canonical H2A around regulatory elements where it facilitates accessibility to the transcriptional machinery (Luk et al., 2010). Hβ-Hα has a very highly charged C-terminal tail with 20 lysines, and we speculate that replacement by the tailless Hζ-Hε variant might be facilitated by lysine acetylation, which would neutralize the basic charge of the Hβ-Hα C-terminal tail and detach it from its electrostatic contact with the acidic DNA wrapping around the core.

Doublet histones are found in some archaeal clades, which led to the proposal that histone doublets made possible the differentiation from homotypic nucleosomes typical of most archaea to heterotypic nucleosomes that later evolved into the four eukaryotic core histones (Malik and Henikoff, 2003). Our evidence that Marseillevirus doublet histones are well-suited for viral packaging is consistent with the possibility that eukaryotic histones have evolved from virus-encoded histone doublets that infected a host proto-eukaryote just as present-day *Marseilleviridae* infect *Acanthamoeba* in oceans around the globe.

There has been considerable recent interest in the hypothesis that the eukaryotic nucleus evolved from a viral factory (Liu and Krupovic, 2022; Talbert and Henikoff, 2021), and if so, the as-yet unknown mechanism whereby a tightly packed viral particle transitions to a fully functional viral factory may shed light on the earliest stages of eukaryotic evolution.

## Methods

### Viruses

*Marseillevirus marseillevirus* Strains T19 (Boyer et al., 2009) and G648 (Sahmi-Bounsiar et al., 2021) were cultured in *Acanthamoeba polyphaga* as described (Pagnier et al., 2013).

### Drosophila cells

Drosophila S2 cells were grown in HyClone SFX Insect Cell Culture Media (Cytiva SH30278.02) supplemented with 18mM L-Glutamine, seeded at 2×10^6^/mL three times per week, and harvested with >95% viability at mid-log phase. A total of 1×10^7^ cells were centrifuged in a swinging-bucket rotor for 4 min at 700xg at 25°C and washed twice in cold 1x PBS. The cell pellet was resuspended in 1 mL TM2+PI (10mM Tris pH 8, 2mM MgCl_2_ + Protease Inhibitor, Sigma 11836170001) and chilled in ice water for 1 min. NP-40 was added to 0.5% and vortexed gently at half maximum speed for ∼3 sec and returned to ice water slurry. Release of nuclei was ascertained by microscopic observation of aliquots until ≥80% of cellular membranes were disrupted (∼3 min). Nuclei were centrifuged 10 min at 150xg at 4°C, washed twice in 1.5 mL TM2+PI and finally resuspended in 200 μL TM2+PI. Each digestion reaction contained either 50K or 150K nuclei per timepoint.

### Viral capsid opening and permeabilization

We followed the viral opening procedure described by Schrad et al., 2020 (Schrad et al., 2020) for Mimivirus, with minor modifications. Each sample from a purified Marseillevirus culture was applied to the membrane of a 7K MWCO Dialysis Unit (Thermo 69562) and dialyzed against 250 mL of 20 mM sodium phosphate buffer adjusted to pH2, 2 mM MgCl_2_, 1 mM PMSF for 2 hr at 25°C, followed by dialysis against 250 mL TM2, 1 mM PMSF for 3 hr. In some experiments viral suspensions were diluted with TM2 prior to dialysis. To equalize the amounts of DNA from Marseillevirus and Drosophila control cells, DNA was extracted from a range of volumes of purified Marseillevirus culture in parallel with a known number of S2 cells. Either 0.5 μL or 1.5 μL of Marseillevirus culture was used per timepoint.

To improve recovery, we included the non-ionic detergent NP-40, which is widely used for chromatin release from cells. Recovery from Marseillevirus T19 particles was vastly improved, ∼85-90% in 0.5% NP-40 using the maximum MNase digestion conditions that had resulted in mostly mononucleosomes in the previous Drosophila experiments. High yields and fragment size distributions were obtained from samples treated with 0.5% NP-40 over a time course ranging from 1 minute to 18 hours. In contrast, 0.1% of the non-ionic detergent digitonin had no effect. As NP-40 permeabilizes membranes by sequestering lipids, whereas digitonin permeabilizes cell membranes by displacing membrane sterols, our results suggest that viral chromatin is enclosed within a lipid membrane that must be permeabilized for MNase to access chromatin.

### MNase-seq

MNase-seq was performed as previously described (Chereji et al., 2019). Briefly, Micrococcal Nuclease (MNase, Sigma N3755) was reconstituted to a concentration of 1 unit per 5 μl with nuclease-free water, aliquoted and stored at -20°C. MNase was thawed on ice and diluted stepwise 1:1 from 1x to 1/512x with TM2 buffer (10 mM Tris pH 8, 2 mM MgCl_2_) where 1x is 20 mU/μL. The volume of each Drosophila or Marseillevirus timepoint was adjusted to 166 μL with TM2+PI (Protease Inhibitor, Sigma 11836170001), and 2.5 μL of each MNase dilution was added with mixing, then heated to 37°C for 1min. MNase was activated by addition of 3.5 μL of 100 mM CaCl_2_ and incubated for 5 min at 37°C. Reactions were stopped with 172 μL of 2xSTOP solution (10mM Tris, 2mM MgCl_2_, 340mM NaCl, 20mM EDTA, 4mM EGTA, 100ug/mL RNase, DNase-free (Sigma11119915001) and DNA was extracted as described below. For reconstituted nucleosomes, we digested with 0.2 U MNase/μg DNA for 7 min at room temperature (Bhardwaj et al., 2020).

We performed digestions over a time course and concentration range that we had previously found to be sufficient for digesting chromatin from Drosophila S2 cells into mono- and oligo-nucleosomes (Chereji et al., 2019). In that study, 1 minute digestions at 2.5 U/million cells had yielded an electrophoretic ‘ladder’ dominated by oligonucleosomes, 5 minute digestions yielded mostly mononucleosomes, and 30 minute digestions resulted in partially degraded mononucleosomes (30 min). We prepared MNase-seq Illumina sequencing libraries from the resulting MNase-digested fragments for both NP-40-treated and untreated libraries, performed paired-end DNA sequencing, and mapped the resulting read pairs to the annotated Marseillevirus T19 genome assembly. On average 97% of fragments mapped to this assembly, confirming the purity of our viral sample. Fragment lengths inferred from sequencing data are smaller than what we observed by Tapestation analysis of purified DNA following MNase digestion (**Figure 1c**), which reflects the selection for smaller fragments during end-polishing and PCR.

### MPE-seq

MPE-seq was performed as described by Ishii et al. (Ishii et al., 2015). Briefly, the opened Marseillevirus and S2 nuclei were treated with hydrogen peroxide across the range of 0.001-1 mM and cleavage was induced by addition of methidiumpropyl-EDTA-Fe(II) (MPE, a generous gift from Jim Kadonaga), which is MPE complexed with ammonium iron(II) sulfate) at 10 μM or 40 μM for 5 min at room temperature. The reaction was quenched with 6 mM bathophenanthroline (Sigma 133159), followed by 2xSTOP buffer (10 mM Tris, 2 mM MgCl_2_, 340 mM NaCl, 20 mM EDTA, 4 mM EGTA, 100 μg/mL RNase, DNase-free) at a volume equal to the sample and total DNA extracted as described below. For reconstituted nucleosomes, we digested with 40 μM MPE using at a ratio of 1 mM H_2_O_2_/650 ng DNA for 5 min at room temperature.

### DNA extraction

Sample volumes were adjusted to ∼340 μL with TM2. To each sample, 3.4 μL 10% SDS and 2.5 μL Proteinase K (20 mg/ml) were added and incubated 30 min 55°C. DNA was extracted once with 350 μL Phenol:Chloroform:Isoamyl Alcohol (25:24:1) in a phase-lock tube (Qiagen cat. no. 129046) and centrifuged 5 min at 16,000xg followed by extraction with 350 μL chloroform. The aqueous layer was transferred to a fresh tube containing 2 μL Glycogen (20 mg/mL Sigma cat. no. 10901393001). DNA was precipitated with three volumes of 100% ethanol and centrifuged at 16,000xg for 30 min at 4^°^C.. The pellet was rinsed twice with 1 ml 80% ethanol, air dried and dissolved in 30 uL 10 mM Tris pH 8.0. Two μL was analyzed using a HDS1000 ScreenTape on an Agilent 4200 TapeStation.

### Proteomics

Viral and reconstituted protein extracts were resolved by 18% SDS-PAGE, and gels were silver-stained using the Pierce kit (Thermo cat. no. 24600). Protein bands were excised, destained, and proteolytically digested as described (Shevchenko et al., 1996). Proteolytic peptides were desalted and analyzed by LC-ESI-MS/MS using a ThermoScientific Orbitrap Fusion mass spectrometer. Mass spectra were analyzed with ThermoScientific Proteome Discoverer version 2.5 software using the *Marseillevirus marseillevirus* translated protein sequences from NCBI (ref. sequence NC_013756.1) appended to the cRAP (https://www.thegpm.org/crap/) common contaminant protein database. Peptide identification results were filtered to a false discovery rate of 1%.

### Library Preparation

Sequencing libraries were prepared from DNA fragments using the KAPA HyperPrep Kit (KAPA cat. no. KK8504) following the manufacturer’s instructions. Libraries were amplified for eight cycles using a 10 sec 60^°^C combined annealing/extension step. Alternatively, a 72^°^C 1 min PCR extension step was added, with no apparent difference in the length distributions. To deplete total DNA samples of large fragments originating from insoluble chromatin prior to library preparation, samples were mixed with ½ volume of HighPrep™ PCR Clean-up System beads (MagBio Genomics cat. no. AC-60005), held 5–10 min, placed on a magnet stand, and the supernatant was collected. Based on the original sample volume, 0.8 volume of beads was added to the supernatant and held 5–10 min. Tubes were placed on a magnet stand, the supernatant discarded and the beads washed 1x with 80% ethanol prior to eluting in 15 uL 10 mM Tris-HCl pH 8. Library size distributions were resolved on an Agilent 4200 TapeStation.

### Expression and purification of Marseillevirus histone tetramers

A pET-24B plasmid containing *Marseillevirus marseillevirus* doublet histones (Hδ-Hγ with a C-terminal 6×His tag and Hβ-Hα) (Valencia-Sanchez et al., 2021) was transformed into *Escherichia coli* BL2-codon plus(DE3)-RIL competent cells (Agilent Technologies) and cultured at 37^°^C in 2xYT-Kanamycin medium. After reaching optical density = 0.6, the cultures were induced with 0.5 mM IPTG at 37^°^C for 3 hr and harvested by centrifugation. Cells were resuspended in 20 mM Tris pH 7.5, 2 M NaCl, 5 mM imidazole, 1 mM β-mercaptoethanol (βme) and lysed on a cell disruptor (AvestinEmulsiflexC3). The extract was clarified by centrifugation, applied to Ni-NTA agarose beads (Qiagen) and eluted with 20 mM Tris pH 7.5, 2 M NaCl, 300 mM imidazole, 1 mM βme. The protein sample was further purified using a Superdex200 26/600 size-exclusion chromatography column (GE Healthcare), and fractions were collected and concentrated to 1.8 mg/ml in 10 mM Tris pH 7.5, 2 M NaCl, 1 mM EDTA, 5 mM βme.

### Purification of Widom 601 DNA array

A plasmid (3_187_widom_601) containing three tandem copies of the Widom 601 nucleosome positioning sequence with a 40-bp linker flanked by the EcoRV restriction site was transformed into DH5α competent cells (ThermoFisher) and cultured in 2xYT-Ampicillin medium overnight. The 3_187_widom_601 DNA insert fragment was excised using EcoRV and purified using a previously published protocol (Dyer et al., 2004).

### Reconstitution of nucleosomes on Widom 601 arrays

*Xenopus laevis* and Marseillevirus nucleosome array reconstitutions were performed as described (Grau et al., 2021). Briefly, nucleosomes arrays were assembled by mixing purified 3_187_widom_601 DNA fragment and histone octamers, followed by overnight gradient salt dialysis in 10 mM Tris–HCl pH 7.5, 1 mM EDTA, 1 mM dithiothreitol (DTT) and 2 M to 0.25 M KCl using a peristaltic pump. Nucleosomes arrays were dialyzed into TCS-50 buffer (20 mM Tris-HCl pH 7.5, 50 mM NaCl, 1 mM EDTA, 1 mM DTT), concentrated, and stored at 4°C until use. Octamer/DNA ratios were optimized using small-scale reactions, and products were verified by native PAGE. The ratio used for Marseillevirus tetramer:DNA was 3.1, and for *X. laevis* octamer:DNA was 3.9.

### Data processing and analysis

Barcoded libraries were mixed to achieve equimolar representation as desired aiming for a final concentration as recommended by the manufacturer for paired-end sequencing on either an Illumina HiSeq 2500 flow cell or a NextSeq flow cell.

For each sample the following analysis steps were performed:

1. Aligned Illumina fastq files to a reference genome with bowtie2 2.4.2 using parameters --end-to-end -- very-sensitive --no-mixed --no-discordant --phred33 -I 10 -X 700
  a. T19: GCF_000887095.1_ViralProj43573_genomic.fna from NCBI Marseillevirus marseillevirus, strain T19 taxon 1559367
  b. G648: G648_26-12-14_Genome_vD.fasta (Supplemental Item 2) ORFs: G648_26-12-14_ORFs_vD.fasta (Supplemental Item 3)
  c. Widom 601 3x Array: Genome_601×3.fasta (Supplemental Item 4)
  d. *Drosophila melanogaster*: dm6 from UCSC. Chronological list of *D. melanogaster* ORFs: https://hgdownload.cse.ucsc.edu/goldenPath/dm6/database/refFlat.txt.gz
2. Extracted aligned fragments from bowtie2 sam files into bed files and divided into four fragment length subgroups: 1-, 1-100, 101-200, 201-300. Steps 3-8 were done for each of the four fragment lengths subgroups.
3. Computed percent GC for 10 base pairs on either end of the mapped fragments.
4. Made normalized count bedgraph files using bedtools 2.30.0 genomecov. Normalized counts are the fraction of counts at each base pair scaled by the size of the reference sequence so that if the scaled counts were uniformly distributed there would be 1 at each position.
5. Obtained annotations and selected ≥ 600 bps long
  a. T19: from NCBI: https://www.ncbi.nlm.nih.gov/nuccore/284504040
  b. G648: Reference (Sahmi-Bounsiar et al., 2021)
  c. *Drosophila melanogaster*: dm6 annotation from UCSC: https://genome.ucsc.edu
6. Extracted normalized count (bedgraph) values for 600 bps inside the selected genes from the TSS and from the TES using deepTools 3.3.1 computeMatrix with these parameters:
  - S <bigwig version of bedgraph file> -R genes.600 --referencePoint TSS -b 0 -a 600
  - S <bigwig version of bedgraph file> -R genes.600 --referencePoint TES -b 600 -a 0
7. Computed autocorrelation for of the extracted values for 1-300 lags using a custom program as R_lag = 1/((n-lag)*var) * sum_1,n-lag(X_t - mean)(X_t+lag - mean) for each gene in each TSS and TES set.
8. Computed the mean and standard deviation of all 299 lags for each gene and reverse ranked the genes by standard deviation, selecting the four genes with the highest standard deviation for each TSS and TES set.

### *De nov*o genome assembly

Paired-end PE50 fragments were trimmed using cutadapt version 2.9 (Martin, 2010) with parameters: cutadapt -j 8 -m 20 --nextseq-trim 20 -a AGATCGGAAGAGCACACGTCTGAACTCCAGTCA -A AGATCGGAAGAGCGTCGTGTAGGGAAAGAGTGT -Z. Trimmed paired-end fastq files were used as input for SPAdes version 3.13.0 (Bankevich et al., 2012) with parameters (−k = kmers): -k 21,27,33,35,37,41. Alignments to Marseillevirus T19 (GenBank NC_013756.1) and G648 (Sahmi-Bounsiar et al., 2021) reference sequences were performed using BLAT (kentinformatics.com).

## Acknowledgments

We thank Jim Kadonaga for MPE, Fred Hutch Genomics Shared Resource for sequencing services, Fred Hutch Proteomics & Metabolomics Shared Resource for mass spectrometry services, and Kami Ahmad for comments on the manuscript. TDB, PBT and SH are supported by the Howard Hughes Medical Institute. PDI, MIV-S and K-JA were supported by a grant from the David and Lucile Packard Foundation.

## Author contributions

Conceptualization: K-JA and SH; Investigation: TDB, PDI, MIV-S, JGH, PBT, BLS, K-JA, SH; Writing – original draft: SH; Writing – review & editing: TDB, PDI, MIV-S, JGH, PBT, K-JA, SH.

---

**Supplementary Table 1:**
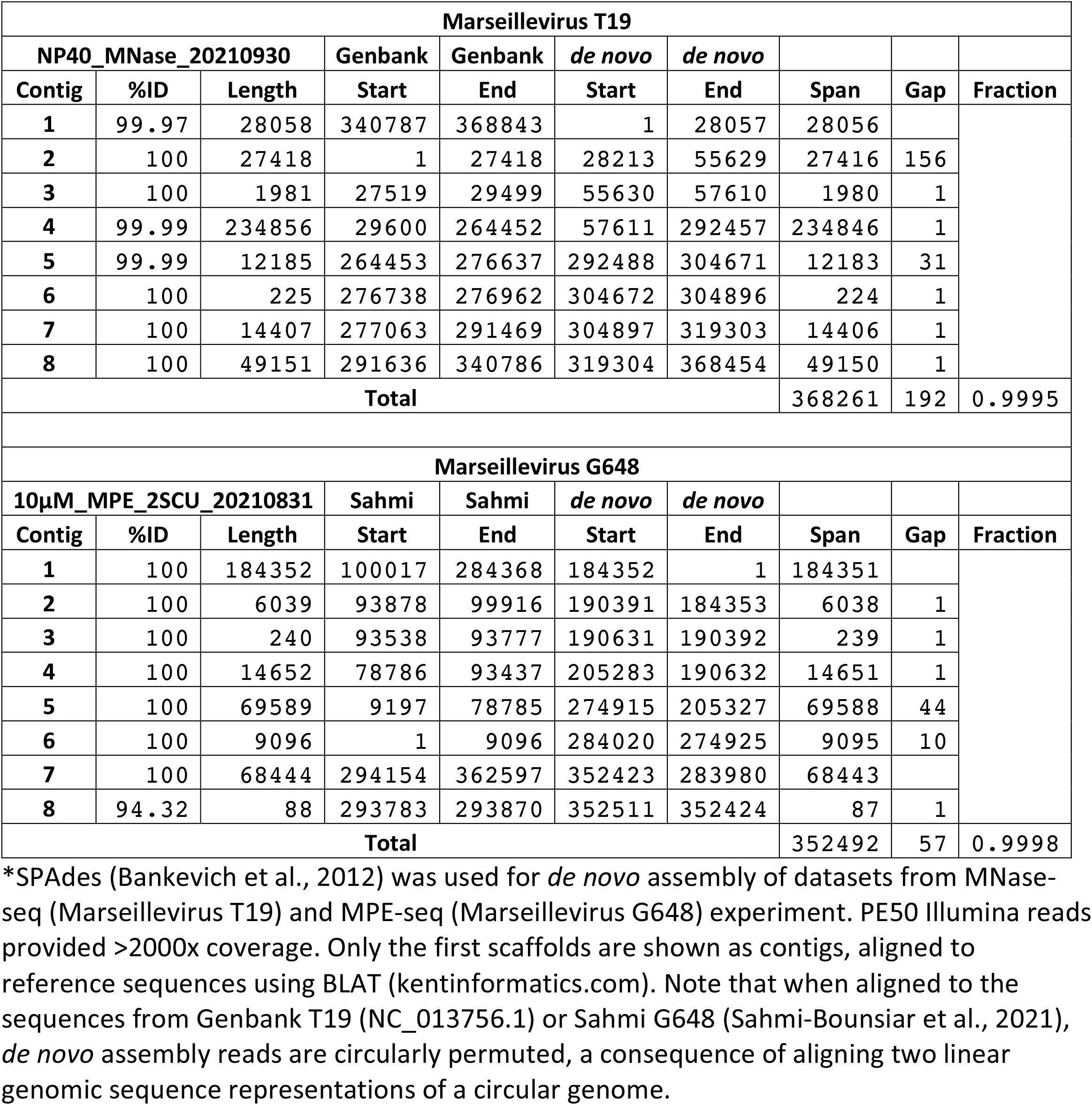
*de novo* assemblies from fragment end reads*.

| Marseillevirus T19 |  |  |  |  |  |  |  |  |  |
| --- | --- | --- | --- | --- | --- | --- | --- | --- | --- |
| NP40_MNase_20210930 |  |  | Genbank | Genbank | *de novo* | *de novo* |  |  |  |
| Contig | %ID | Length | Start | End | Start | End | Span | Gap | Fraction |
| 1 | 99.97 | 28058 | 340787 | 368843 | 1 | 28057 | 28056 |  |  |
| 2 | 100 | 27418 | 1 | 27418 | 28213 | 55629 | 27416 | 156 |  |
| 3 | 100 | 1981 | 27519 | 29499 | 55630 | 57610 | 1980 | 1 |  |
| 4 | 99.99 | 234856 | 29600 | 264452 | 57611 | 292457 | 234846 | 1 |  |
| 5 | 99.99 | 12185 | 264453 | 276637 | 292488 | 304671 | 12183 | 31 |  |
| 6 | 100 | 225 | 276738 | 276962 | 304672 | 304896 | 224 | 1 |  |
| 7 | 100 | 14407 | 277063 | 291469 | 304897 | 319303 | 14406 | 1 |  |
| 8 | 100 | 49151 | 291636 | 340786 | 319304 | 368454 | 49150 | 1 |  |
| Total |  |  |  |  |  |  | 368261 | 192 | 0.9995 |
| Marseillevirus G648 |  |  |  |  |  |  |  |  |  |
| 10μM_MPE_2SCU_20210831 |  |  | Sahmi | Sahmi | *de novo* | *de novo* |  |  |  |
| Contig | %ID | Length | Start | End | Start | End | Span | Gap | Fraction |
| 1 | 100 | 184352 | 100017 | 284368 | 184352 | 1 | 184351 |  |  |
| 2 | 100 | 6039 | 93878 | 99916 | 190391 | 184353 | 6038 | 1 |  |
| 3 | 100 | 240 | 93538 | 93777 | 190631 | 190392 | 239 | 1 |  |
| 4 | 100 | 14652 | 78786 | 93437 | 205283 | 190632 | 14651 | 1 |  |
| 5 | 100 | 69589 | 9197 | 78785 | 274915 | 205327 | 69588 | 44 |  |
| 6 | 100 | 9096 | 1 | 9096 | 284020 | 274925 | 9095 | 10 |  |
| 7 | 100 | 68444 | 294154 | 362597 | 352423 | 283980 | 68443 |  |  |
| 8 | 94.32 | 88 | 293783 | 293870 | 352511 | 352424 | 87 | 1 |  |
| Total |  |  |  |  |  |  | 352492 | 57 | 0.9998 |

\*SPAdes (Bankevich et al., 2012) was used for *de novo* assembly of datasets from MNase-seq (Marseillevirus T19) and MPE-seq (Marseillevirus G648) experiment. PE50 Illumina reads provided >2000x coverage. Only the first scaffolds are shown as contigs, aligned to reference sequences using BLAT (kentinformatics.com). Note that when aligned to the sequences from Genbank T19 (NC\_013756.1) or Sahmi G648 (Sahmi-Bounsiar et al., 2021), *de novo* assembly reads are circularly permuted, a consequence of aligning two linear genomic sequence representations of a circular genome.

**Supplementary Figure 1:**
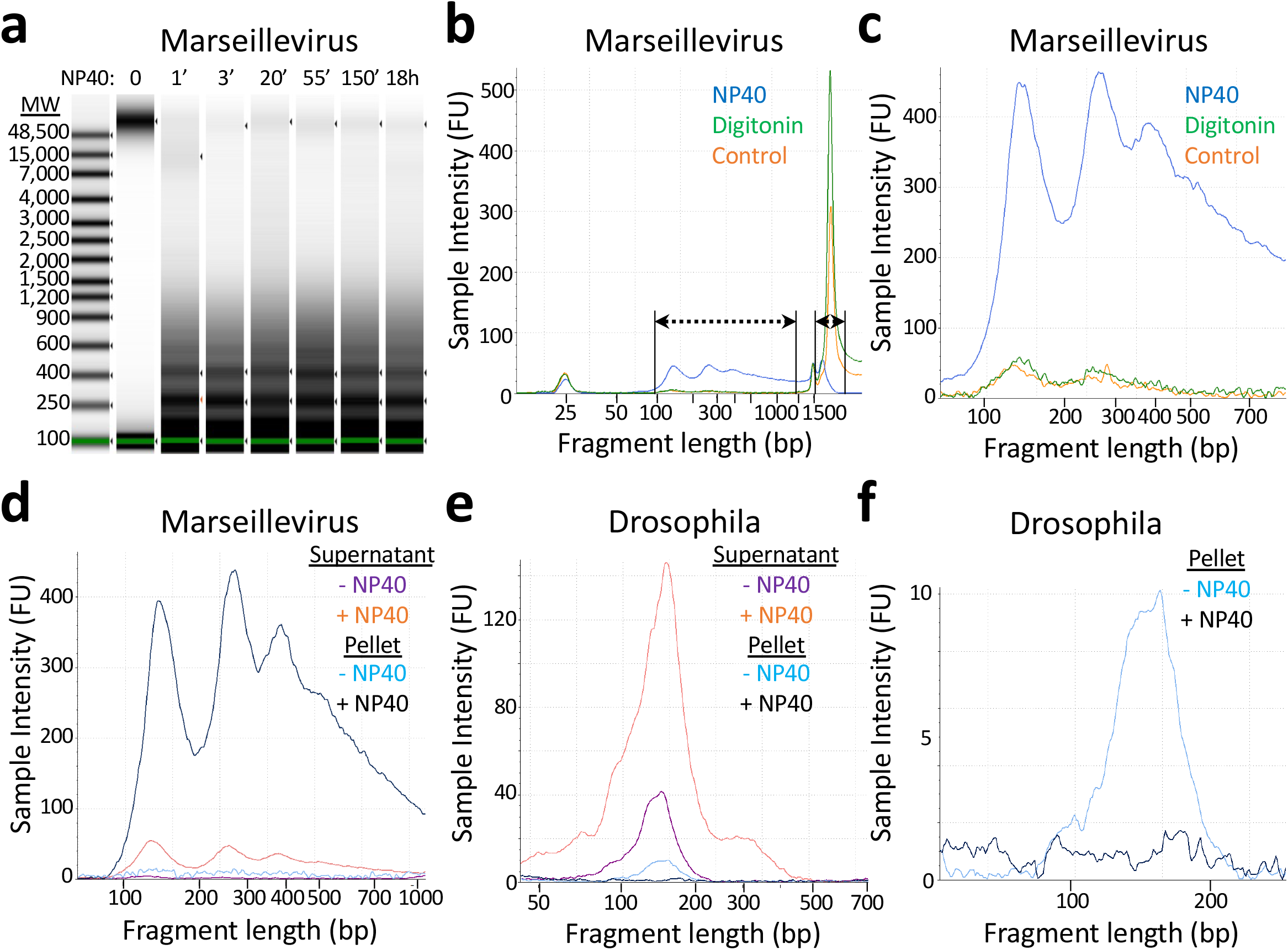
Marseillevirus nucleosomes are mostly insoluble after MNase treatment. **a)** Tapestation gel image of DNA purified from breached Marseillevirus T19 virus particles after NP40 treatment for times as indicated and 5 min MNase digestion. (**b**) Profiles of Marseillevirus fragment lengths showing the ranges of signal used for calculating recovery: 100-1000 bp for MNase-released chromatin and >2 kb for undigested DNA. (**c**) Zoom-in of (b) showing relative recoveries. (**d**) Profiles of Marseillevirus T19 fragment lengths ± NP-40 after centrifugation to separate supernatant and pellet, showing that even after NP40 treatment, the large majority of Marseillevirus chromatin remains insoluble after MNase digestion. (**e**) Same as (d) except for Drosophila, showing that NP40 + MNase quantitatively solubilizes chromatin. (**f**) Zoom-in of the Drosophila pellet fractions (e) showing that no detectable DNA remains after NP40 extraction.

**Supplementary Figure 2:**
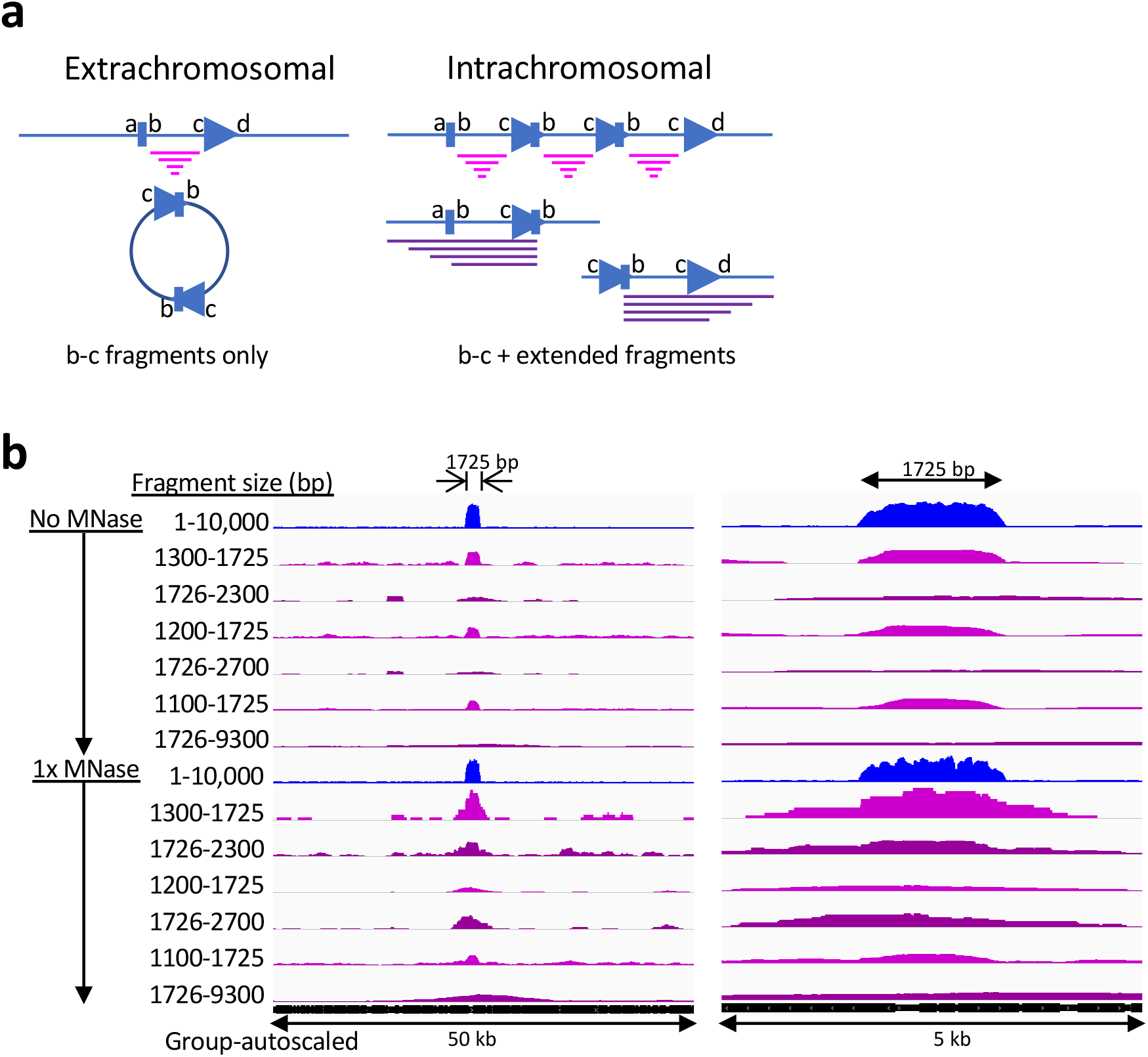
Regional over-representation of DNA is intrachromosomal. (**a**) Both extrachromosomal and intrachromosomal tandem repeat models predict novel c/b junctions and mappable fragments (magenta) above background that are shorter than the over-represented region, but only the intrachromosomal model predicts extended fragments above background that start upstream of point a and span downstream beyond point b and start downstream of point d and span upstream beyond point c. (**b**) Tracks show the region around the 1725-bp over-represented region observed in No-MNase- and MNase-seq data for the indicated fragment size intervals. The enrichment of 1725-bp fragments spanning point b on the left and point c on the right relative to background is seen for all comparisons, as predicted for the intrachromosomal model, but not the extrachromosomal model.

**Supplementary Figure 3:**
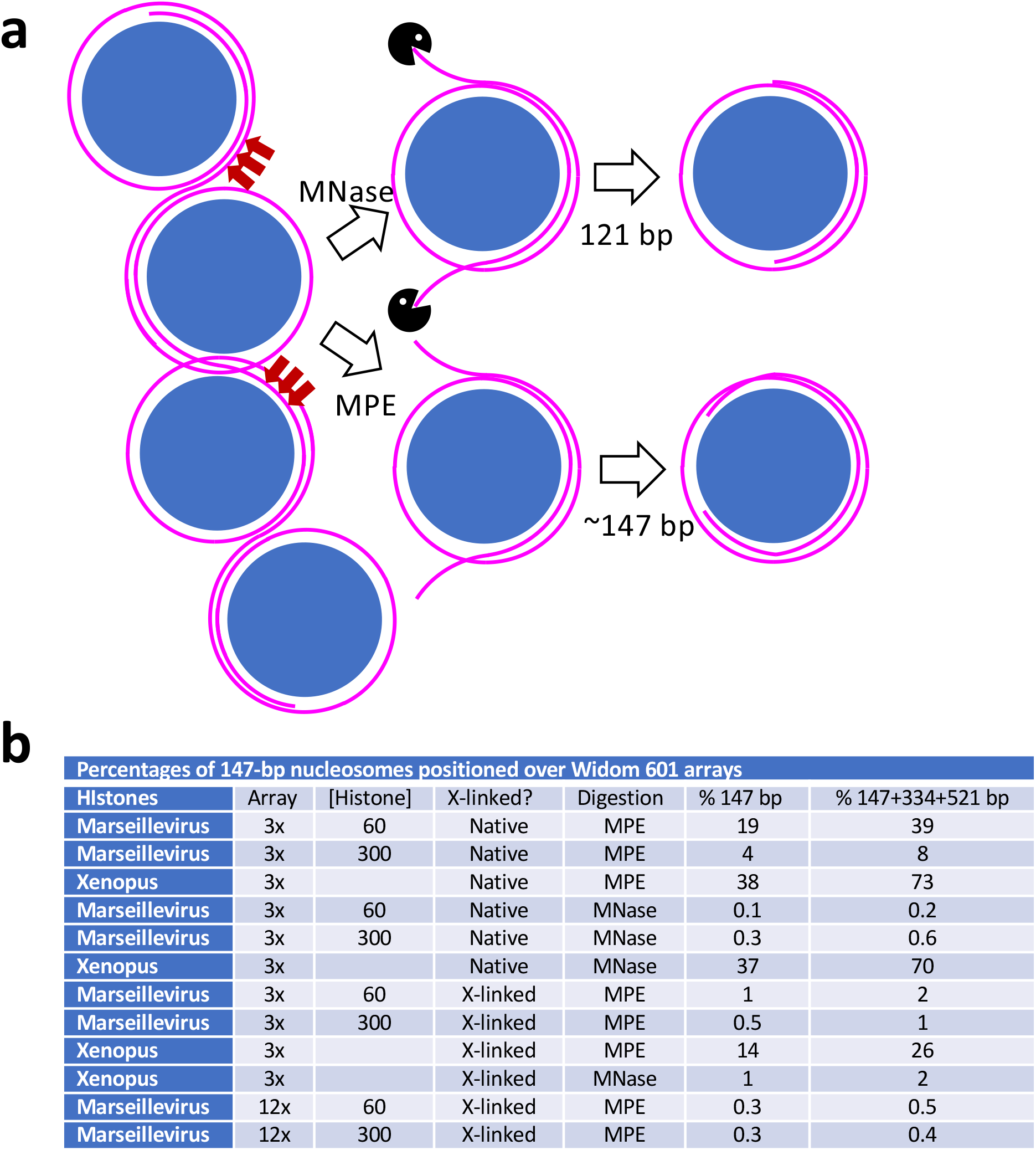
Model for MNase and MPE cleavage patterns. (**a**) Closely abutted and tightly wrapped nucleosomes with ∼1.4 wraps (121 bp) are almost completely inaccessible to MNase and MPE, with partial accessibility preferentially but not exclusively where the wrap shifts from one nucleosome to the next (red arrows). Double-strand cleavages on neighboring nucleosomes release the particle in between, leaving unwrapped ends. Because of its exonuclease activity MNase “nibbles” on the ends down to 121 bp, and when dinucleosomes are released, nibbling results in two adjacent 121-bp particles, and so on. Cleavages within adjacent particles separated by a single DNA wrap (not shown) will release fragments of variable size averaging ∼70 bp. Upon nucleosome release by MPE the unwrapped DNA ends immediately snap onto the exposed basic surface of the histone core resulting in a broad distribution of fragment sizes with a peak at ∼147 bp. (**b**) Percentages of precisely positioned 147-bp nucleosomes (%147 bp) and cleavages (%147+334+521 bp).

**Supplementary Figure 4:**
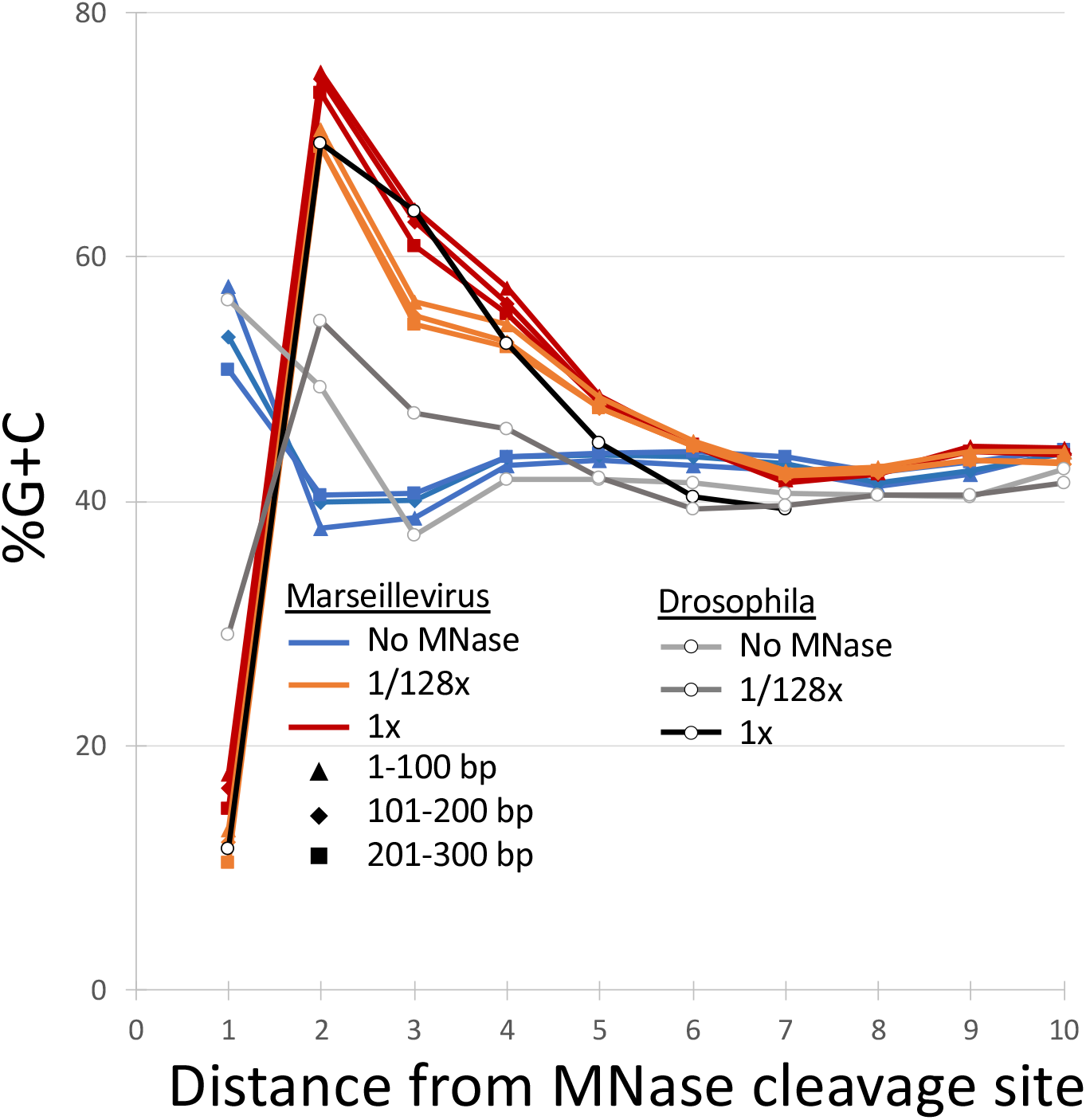
Preferential cleavage between A/T base pairs in Marseillevirus and Drosophila. MNase is an endo-exonuclease that cleaves preferentially at A/T-rich DNA then ‘nibbles’ on ends until it reaches a G/C-rich ‘clamp’, as observed in our MNase-seq data for both Marseillevirus and Drosophila, whose genomes are both 44-45% G+C overall.

**Supplementary Figure 5:**
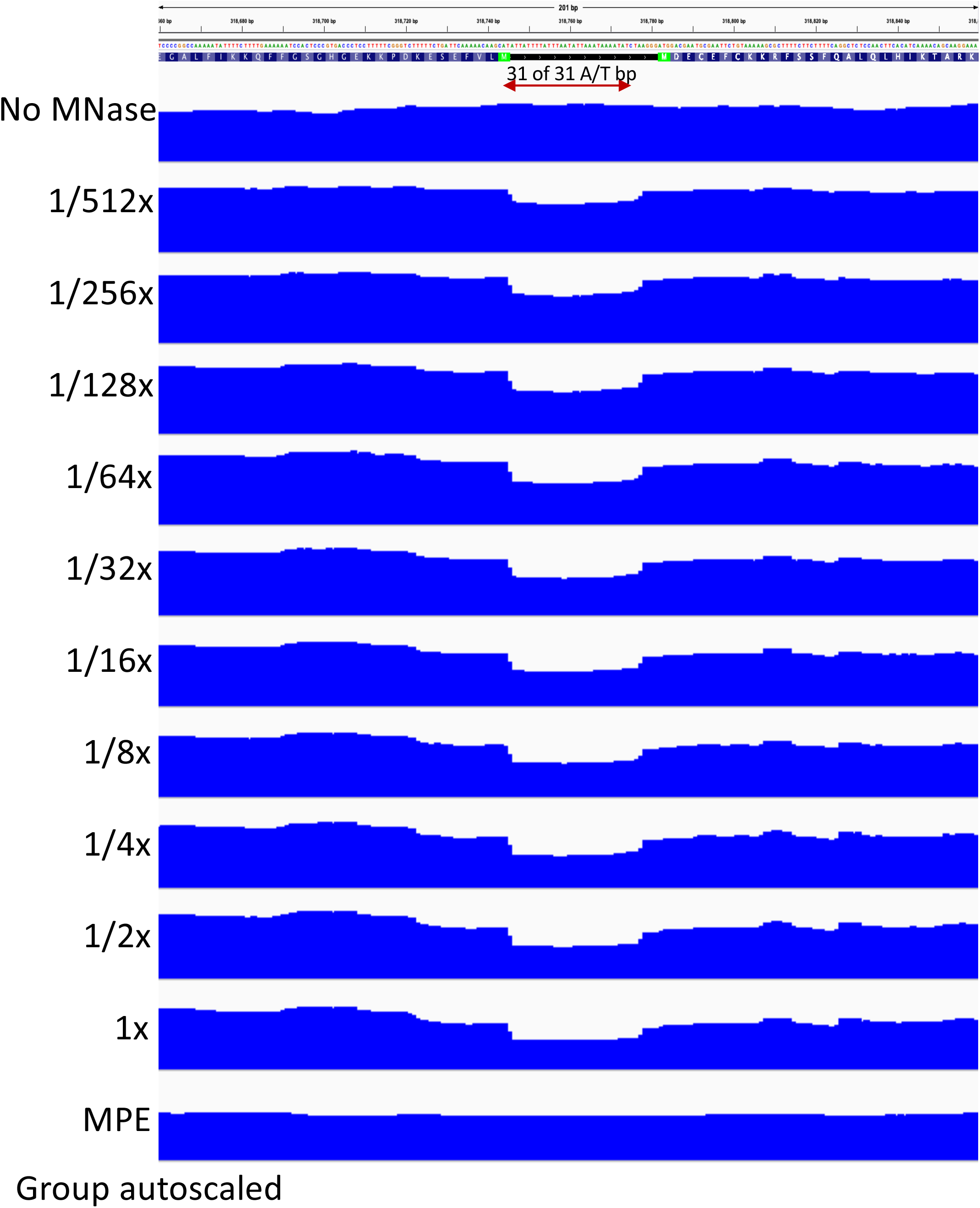
Preferential cleavage between A/T base pairs in Marseillevirus. Track names refer to the 1x-512x MNase series and the 1000 nM sample track in Figure 3a.

**Supplementary Figure 6:**
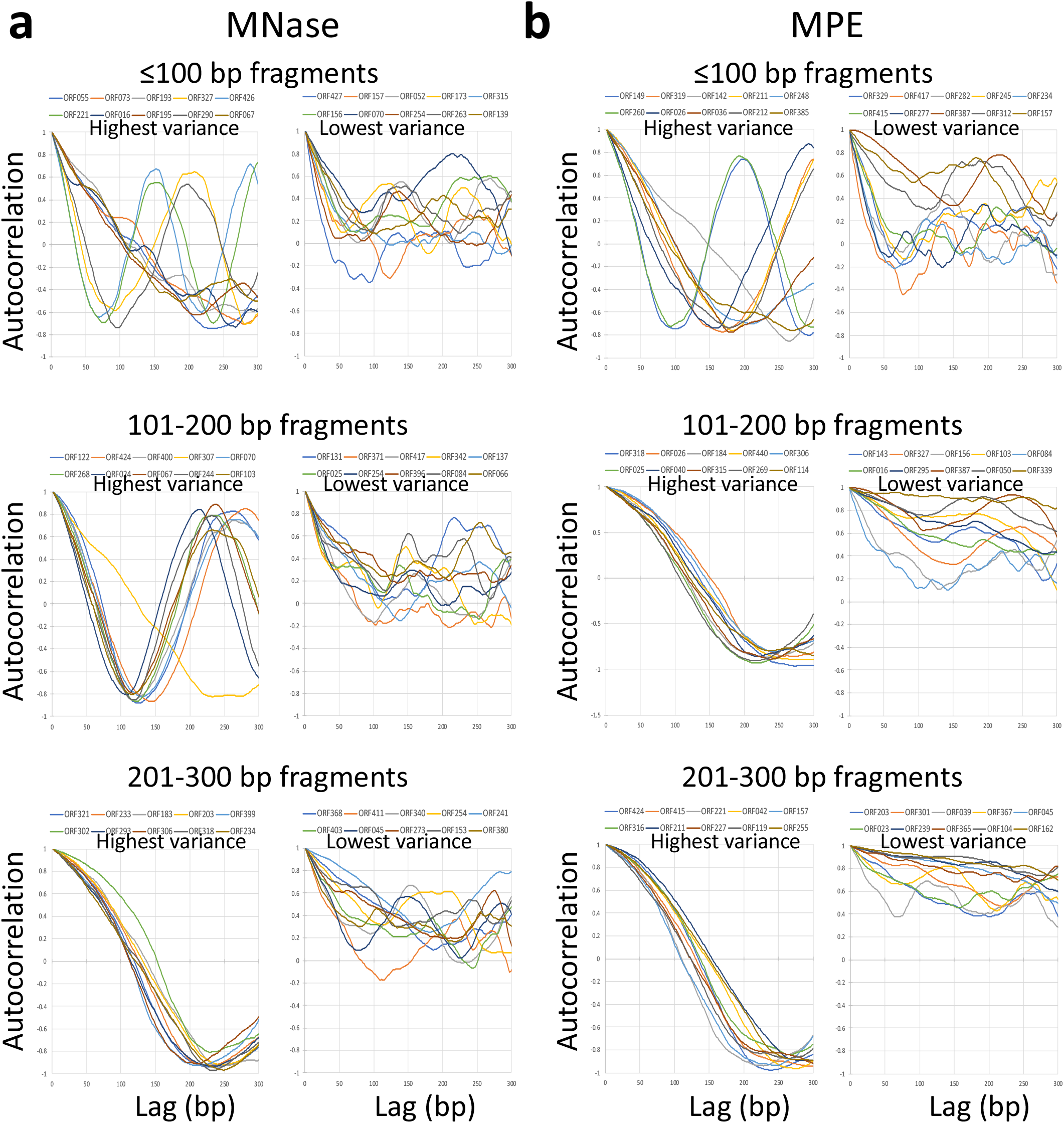
Autocorrelation illustrates periodicities over individual Marseillevirus gene bodies. To sensitively detect genes most likely to be phased, we plotted autocorrelations over the 600-bp span of each ORF for the 101-200 bp fragment size class using MNase-seq and MPE-seq data. Autocorrelations (−1 to +1) in 1-bp lag steps over a 5’-aligned 300-bp span are plotted for each of the 10 ORFs ≥600 bp with the highest and lowest variance in amplitude for each size class for (**a**) MNase-seq and (**b**) MPE-seq.

